# A compact glutamic acid decarboxylase 67 promoter enables inhibitory neuron-targeted AAV gene therapy for treatment-resistant epilepsy

**DOI:** 10.64898/2025.12.07.692482

**Authors:** Yuuki Fukai, Ayumu Konno, Nobutake Hosoi, Karin Miyakawa, Ryosuke Kaneko, Hirokazu Hirai

## Abstract

Epilepsy often becomes treatment-resistant, partly due to impaired inhibitory neurotransmission and reduced γ-aminobutyric acid (GABA) function. Enhancing inhibitory neuron activity via gene therapy may restore excitation–inhibition (E/I) balance. We developed a compact 410-bp glutamic acid decarboxylase 67 promoter (cmGAD67) that enables strong, selective transgene expression in inhibitory neurons while preserving AAV packaging capacity. When delivered systemically, AAV vectors carrying cmGAD67 preferentially targeted parvalbumin interneurons and supported effective circuit manipulation.

To evaluate therapeutic potential, we expressed glutamic acid decarboxylase 65 (GAD65) under cmGAD67 (AAV-GAD65) in pentylenetetrazole (PTZ) epilepsy models. Systemic AAV-GAD65 suppressed abnormal delta oscillations, reduced seizure-like events, normalized anxiety-like behavior, and improved survival in a severe PTZ paradigm. Biochemical analyses confirmed increased cortical and hippocampal GABA levels, linking behavioral and electrophysiological improvements to enhanced inhibitory neurotransmitter synthesis.

Prior clinical evidence indicates that AAV-GAD65 delivery to the subthalamic nucleus is safe and effective in Parkinson’s disease. Building on this foundation, our findings establish the cmGAD67 promoter as a powerful platform for inhibitory neuron-targeted AAV gene therapy and highlight AAV-cmGAD67-GAD65 as a promising approach for treatment-resistant epilepsy and other disorders involving disrupted E/I balance.

## INTRODUCTION

The maintenance of excitation–inhibition (E/I) balance is essential for higher brain functions such as cognition, behavior, and learning. In the cerebral cortex, the activity of excitatory pyramidal neurons is tightly regulated by inhibitory interneurons. Impairment of inhibitory neuron function shifts the E/I balance toward excitation, a pathological state implicated in the development of diverse neuropsychiatric disorders, including epilepsy, anxiety disorders, autism spectrum disorders, and schizophrenia ^1–3^. Thus, strategies that specifically modulate inhibitory neuron function hold promise for the treatment of these conditions.

GABAergic interneurons in the cortex and hippocampus release γ-aminobutyric acid (GABA), the principal inhibitory neurotransmitter, which suppresses excitatory signaling through activation of postsynaptic GABA receptors. GABA is synthesized from glutamate by glutamic acid decarboxylase (GAD), an enzyme expressed selectively in inhibitory neurons. Two isoforms exist, GAD65 (encoded by Gad2) and GAD67 (encoded by Gad1) ^4^. GAD65 is localized mainly to synaptic vesicles and contributes to activity-dependent GABA synthesis, whereas GAD67 is distributed throughout the cytoplasm and provides a steady supply of GABA critical for network stability ^4–6^. Consistent with these roles, GAD65-deficient mice are viable, whereas GAD67-deficient mice die shortly after birth ^7,8^, underscoring the essential and constitutively strong activity of the GAD67 promoter.

Previously, we identified a 2.5-kb fragment upstream of exon 1 of the Gad2 gene that functions as an inhibitory neuron-specific promoter, designated mGAD65(delE1) ^9^. Although this promoter can be incorporated into AAV vectors (Addgene #177316), its large size occupies more than half of the limited AAV packaging capacity (∼4.7 kb), and its weak activity limits the expression of therapeutic genes, particularly after systemic delivery.

To overcome these limitations, we conducted a systematic analysis of the Gad1 locus and identified a compact 410-bp sequence, termed the compact mouse GAD67 (cmGAD67) promoter. Despite its short length, the cmGAD67 promoter exhibits strong activity and high inhibitory neuron specificity when packaged into AAV vectors. In this study, we used AAV vectors equipped with the cmGAD67 promoter to manipulate inhibitory neuron function, assess the resulting effects on brain activity and behavior, and evaluate therapeutic outcomes in epilepsy models. Importantly, the translational feasibility of GAD65-based therapy has already been demonstrated in humans: a first-in-human clinical trial reported that AAV-GAD65 delivery to the subthalamic nucleus was safe and effective in patients with Parkinson’s disease ^10^. Building on this precedent, we highlight the potential of cmGAD67-driven AAV-GAD65 as a novel gene therapy strategy for treatment-resistant epilepsy.

## RESULTS

### Development of a compact GAD67 promoter with strong activity and inhibitory neuron specificity

Inhibitory neuron-specific promoters are essential for AAV-based gene delivery to restore E/I balance in epilepsy and related disorders. Previously, we identified an upstream fragment of the GAD65 gene [mGAD65(delE1)] ^9^, which confers inhibitory neuron specificity when packaged into AAV vectors. However, this promoter is relatively large (2.5 kb) and drives weak expression, occupying a substantial fraction of the limited AAV packaging capacity (∼4.7 kb) and thereby restricting its translational utility.

To overcome these constraints, we performed a systematic analysis of the Gad1 locus and identified a minimal 410-bp regulatory sequence, designated the compact mouse GAD67 (cmGAD67) promoter (Addgene plasmid #245935). Despite its short length, cmGAD67 exhibited strong transcriptional activity and high inhibitory neuron specificity when incorporated into AAV vectors and delivered systemically (Figure 1). Detailed deletion mapping and promoter dissection are shown in supplementary figures S1–S6.

**Figure 1.**
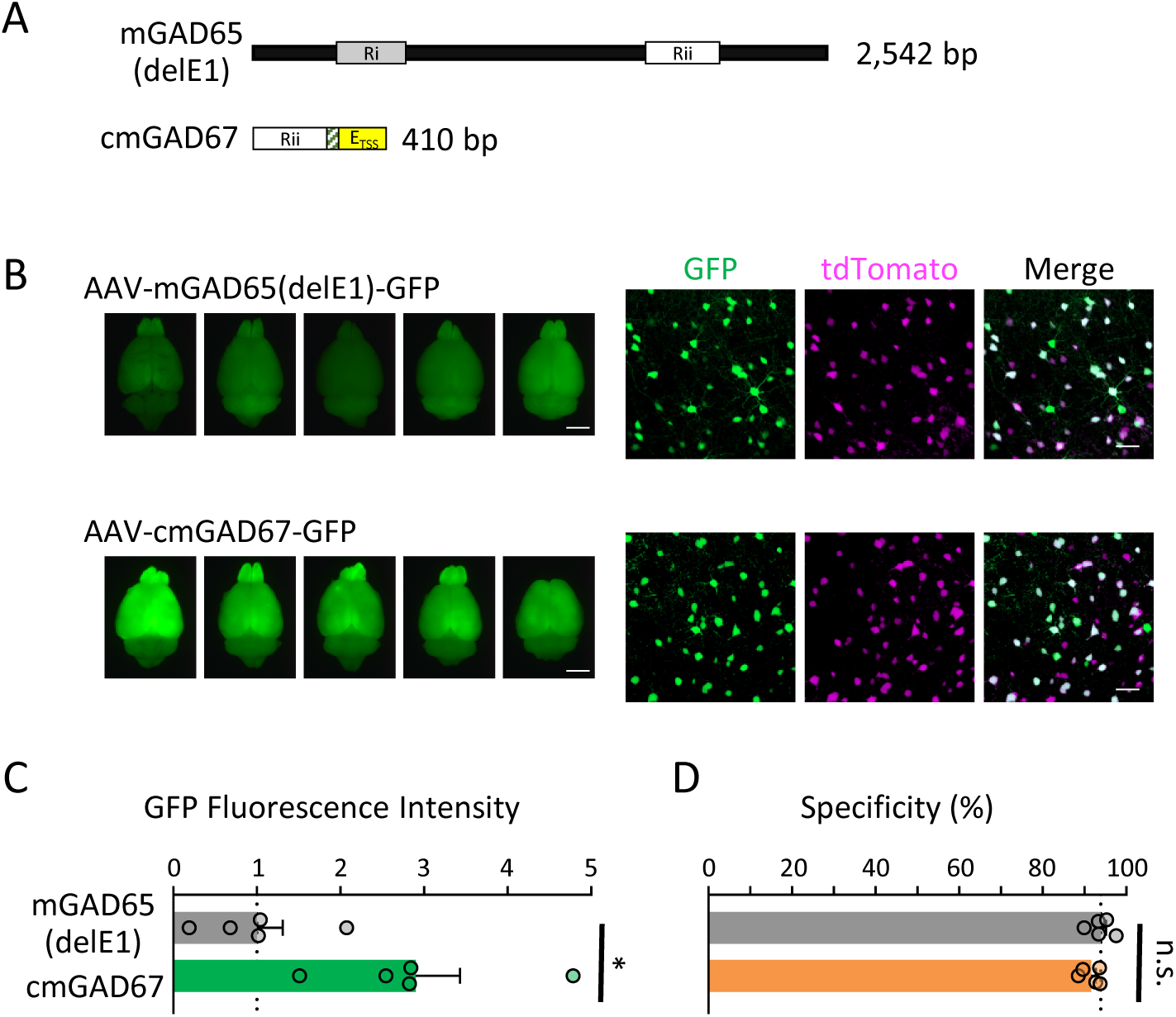
Characterization of the compact GAD67 promoter (cmGAD67) (A) Genomic organization of the conventional mGAD65(delE1) promoter and the compact GAD67 promoter (cmGAD67; 410 bp) at the same scale. The cmGAD67 promoter comprises a minimal regulatory region containing the E_TSS_ core promoter and a downstream enhancer (Rii). Detailed mapping of Ri, Rii, and E_TSS_ subregions is provided in supplementary figures S1–S3. (B) Representative fluorescence images of whole brains and cortical sections from VGAT-tdTomato mice intravenously injected with AAV-PHP.eB expressing GFP under the control of the mGAD65(delE1) or cmGAD67 promoter. GFP (green) expression is largely restricted to tdTomato-positive inhibitory neurons (magenta). Scale bars: whole brain, 5 mm; cortical sections, 50 μm. (C) Quantification of whole-brain GFP fluorescence intensity. Fluorescence intensity in the control mGAD65(delE1) group was normalized to 1. (D) Proportion of GFP-positive cells co-expressing tdTomato, reflecting inhibitory neuron specificity. Data are presented as mean ± SEM (n = 5 mice). Statistical significance was determined using an unpaired t-test; *p < 0.05, n.s., not significant.

### Preferential targeting of PV-positive interneurons by the cmGAD67 promoter

To further characterize the neuronal subtypes transduced by the cmGAD67 promoter, we examined its expression in parvalbumin (PV)- and somatostatin (SST)-positive (PV^+^ and SST^+^) interneurons, which represent two major classes of cortical inhibitory neurons. Intravenous administration of AAV-PHP.eB-cmGAD67-GFP resulted in robust GFP expression predominantly in PV^+^ interneurons, with a smaller fraction of SST^+^ interneurons labeled (Figure 2A-C). Quantitative analyses showed that ∼85% of PV^+^ neurons expressed GFP, whereas fewer SST^+^ neurons, approximately 50%, were transduced (Figure 2D). When examining the distribution of GFP-positive cells, the majority were PV^+^ interneurons, with smaller contributions from SST^+^ and other inhibitory subtypes (Figure 2E).

**Figure 2.**
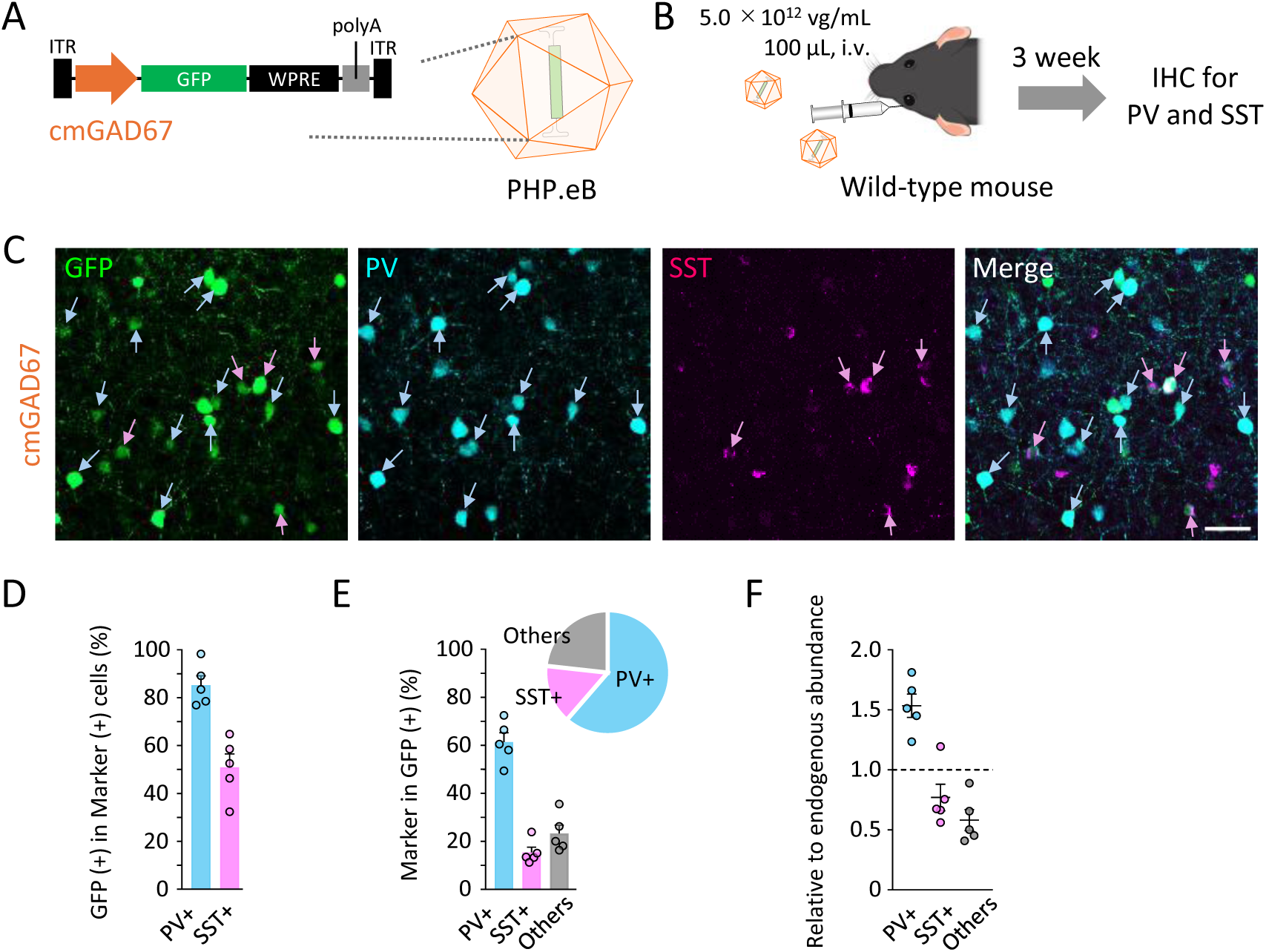
Preferential expression of the cmGAD67 promoter-driven AAV in PV-positive interneurons, with limited targeting of SST-positive interneurons. (A) Schematic representation of the AAV-PHP.eB vector expressing GFP under the control of the cmGAD67 promoter. (B) Experimental design. Wild-type mice were intravenously injected with AAV-PHP.eB-cmGAD67-GFP (hereafter AAV-GFP; 5.0 × 10¹² vg/mL, 100 μL), and brains were analyzed 3 weeks later by immunohistochemistry for parvalbumin (PV) and somatostatin (SST). (C) Immunohistochemical staining of the motor cortex for PV (blue channel) and SST (magenta channel) in mice injected with PHP.eB carrying the cmGAD65 promoter. Cyan arrows indicate PV-positive (PV^+^) neurons, and magenta arrows indicate SST-positive (SST^+^) neurons. Scale bars, 50⍰μm. (D) Quantification of the proportion of PV^+^ or SST^+^ interneurons that express GFP, demonstrating preferential targeting of PV^+^ interneurons. (E) Bar graph and pie chart showing the composition of PV^+^, SST^+^, and double-negative cells among GFP-expressing neurons, showing that the majority of GFP-positive neurons are PV^+^ interneurons, with smaller contributions from SST^+^ and other inhibitory populations. Data represent mean ± SEM (n = 5 mice). (F) Relative tropism of the cmGAD67 promoter-driven AAV-PHP.eB vectors for different GABAergic neuron subtypes (PV⁺, SST⁺, or others) in the mouse cortex. In the mouse cortex, PV⁺ neurons account for 40%, SST⁺ neurons for 15%, and other subtypes for 45% of all GABAergic neurons ^9^. The ratio of each GFP-expressing subtype (from E) was divided by its expected proportion in the cortex to calculate relative enrichment. A value near 1 indicates no subtype preference, whereas a value greater than 1.5 for PV⁺ neurons indicates selective gene expression in this subtype.

In our previous study, GABAergic neuron subtypes in the mouse motor cortex consisted of 40% PV⁺, 15% SST⁺, and 45% other interneurons ^9^. The ratio of each GFP-expressing subtype was normalized to the proportion in the cortex to calculate relative enrichment. A value close to 1 indicates no subtype preference, whereas a value greater than 1.5 for PV⁺ neurons indicates that the cmGAD67 promoter drives preferential gene expression in this subtype (Figure 2F).

These results demonstrate that the cmGAD67 promoter preferentially drives transgene expression in PV^+^ interneurons, a property that is particularly advantageous for manipulating excitatory output through perisomatic inhibition ^11,12^. Additional comparisons with the cmGAD65 promoter are provided in supplementary figure 7.

### Application of the cmGAD67 promoter for optogenetic and chemogenetic manipulation

We next examined whether the cmGAD67 promoter could be used to bidirectionally manipulate inhibitory neuron function through optogenetics and chemogenetics. For neuronal activation, we employed GFP-fused channelrhodopsin-2 (ChR2), and for inhibition, GFP-fused Gi-DREADD was used (Figure. 3A). AAV-PHP.eB-cmGAD67-ChR2 was intravenously administered to VGAT-tdTomato mice ^13^, and patch-clamp recordings were performed in cortical slices three weeks later. In contrast, AAV-PHP.eB-cmGAD67-hM4D(Gi) (Gi-DREADD) was injected bilaterally into the hippocampus of wild-type mice, and three weeks later, behavioral seizures and cortical electrocorticogram (ECoG) recordings from the motor cortex were assessed.

**Figure 3.**
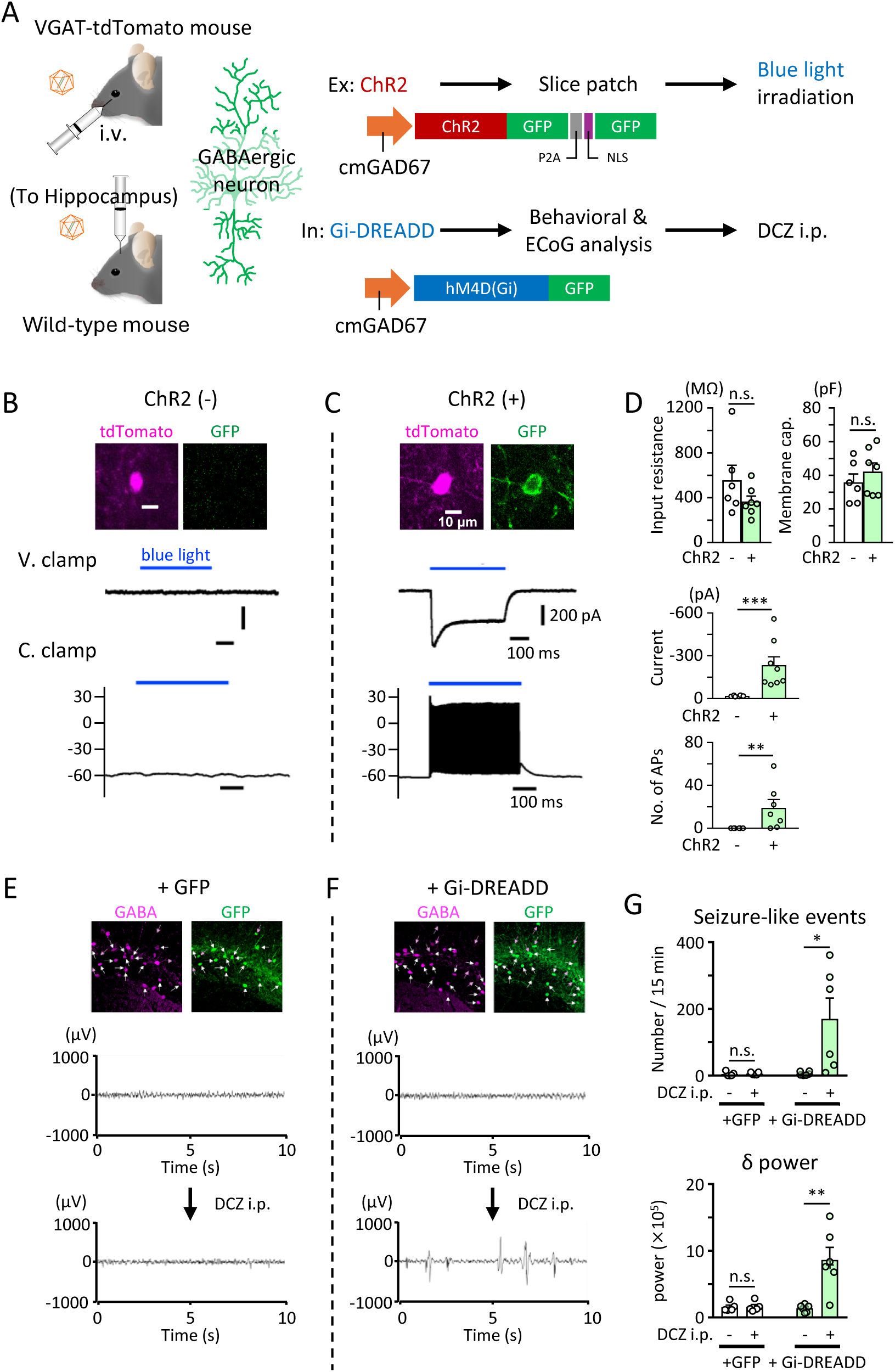
Application of the cmGAD67 promoter for optogenetic and chemogenetic manipulation of inhibitory neurons. (A) Schematic diagrams of AAV constructs expressing channelrhodopsin-2 (ChR2) or the inhibitory designer receptor hM4Di (Gi-DREADD) under the control of the cmGAD67 promoter. (B, C) Representative recordings from cortical tdTomato-positive inhibitory neurons without (B) or with (C) ChR2 expression under voltage-clamp (V. clamp) and current-clamp (C. clamp) conditions, showing robust light-evoked responses only in ChR2(+) inhibitory neurons. (D) Pooled data demonstrate no change in passive membrane properties but significant increases in light-evoked inward current and action potential (AP) firing in inhibitory neurons following optical stimulation (n = 6-8 mice; **p<0.01, ***p < 0.001, Mann-Whitney test; mean ± SEM). (E, F) Representative ECoG traces from mice injected with AAV-cmGAD67-GFP or AAV-cmGAD67-hM4Di-GFP, showing increased network activity only in Gi-DREADD–expressing mice following systemic administration of DCZ. (G) Quantification of ECoG traces indicate a significant increase in the number of abnormal spikes (epileptiform discharges) and in delta power after chemogenetic inhibition (n = 5-6 mice; *p < 0.05, **p < 0.01, paired t-test; mean ± SEM).

In tdTomato-positive inhibitory neurons lacking GFP labeling (i.e., without ChR2 expression), blue light stimulation induced neither photocurrents in voltage-clamp mode nor membrane potential changes in current-clamp mode (Figure. 3B). By contrast, ChR2-expressing tdTomato-positive inhibitory neurons exhibited inward photocurrents in response to blue light under voltage clamp (Figure 3C) and robust action potential bursts under current clamp (Figure. 3C). Quantitative analysis is summarized in Figure 3D. Input resistance and membrane capacitance of inhibitory neurons did not differ between ChR2-expressing and non-expressing cells.

For chemogenetic manipulation, GFP expression from AAV-PHP.eB-cmGAD67-hM4D(Gi) was localized to GABA-positive inhibitory neurons, as confirmed by immunostaining (Figures 3E and F, top). In control mice injected with AAV-PHP.eB-cmGAD67-GFP, systemic administration of deschloroclozapine (DCZ) produced no behavioral or electrophysiological effects (Supplementary movie 1 and Figure 3E). In contrast, Gi-DREADD–expressing mice exhibited marked behavioral seizures (Supplementary movie 2). Correspondingly, ECoG recordings revealed epileptiform activity characterized by abnormal spike-and-wave discharges after DCZ administration in Gi-DREADD mice (Figure 3F). Quantification of abnormal spikes (epileptic discharges) and delta power during 10–25 min after DCZ administration demonstrated significant increases in the Gi-DREADD group but not in GFP controls (Figure 3G).

### AAV-GAD65 suppresses epileptiform activity in the PTZ kindling model

We next examined the effects of AAV-cmGAD67-GAD65 (hereafter AAV-GAD65) on epileptiform activity in the PTZ kindling model. Repeated intraperitoneal injections of PTZ (30 mg/kg) resulted in progressive increases in seizure severity as assessed by the Racine scale (Figures 4A and 4B), confirming successful induction of chronic epilepsy. ECoG recordings obtained 1 and 22 days after the final PTZ injection (Figure 4C) revealed persistent epileptiform discharges, characterized by frequent seizure-like events and, although slightly reduced, a sustained increase in delta power in PTZ-treated control mice (Figure 4D).

**Figure 4.**
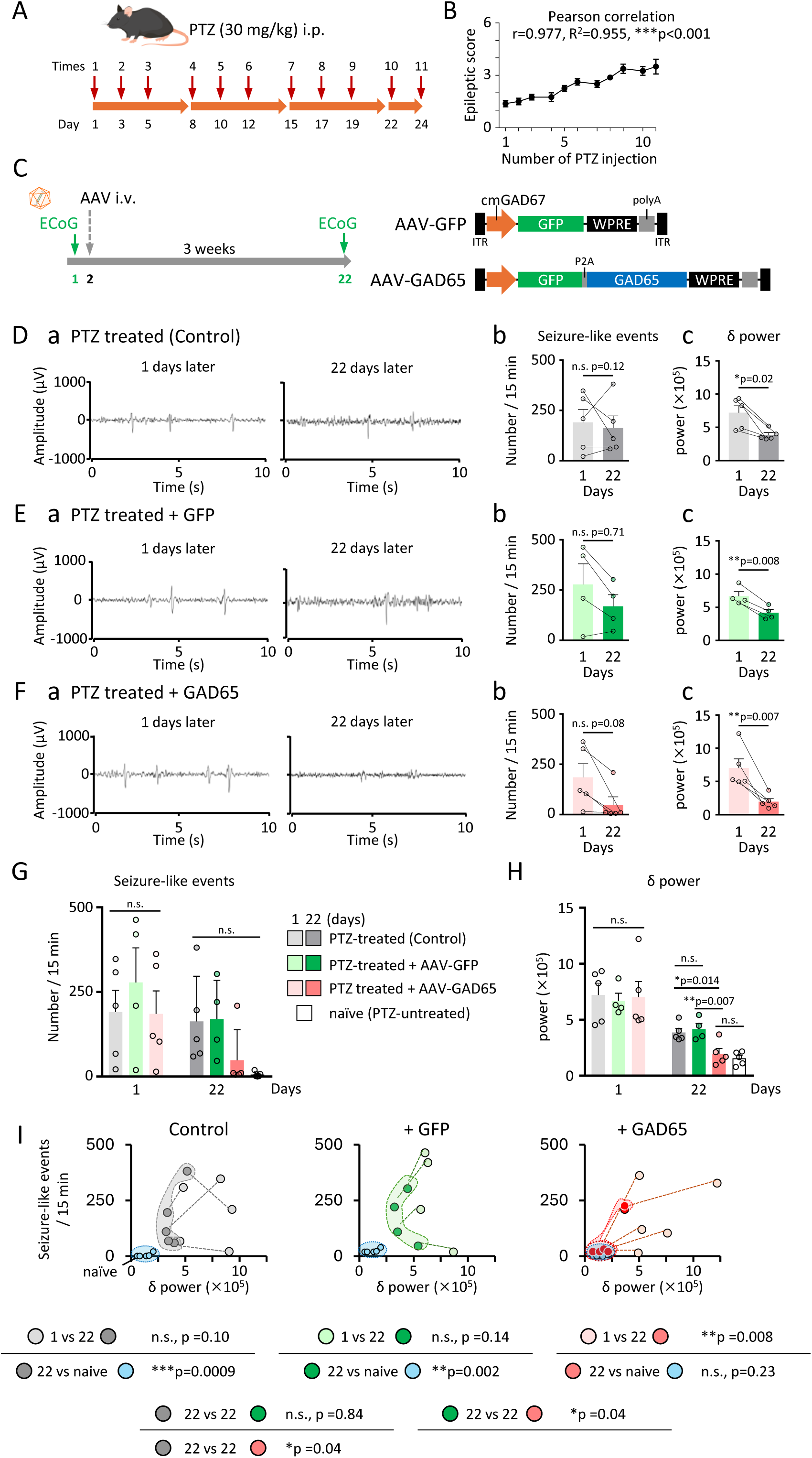
Electrophysiological assessment of PTZ-induced epilepsy and effects of AAV-cmGAD67-GAD65 (hereafter AAV-GAD65) (A) Experimental timeline of PTZ kindling. Mice received intraperitoneal injections of PTZ (30 mg/kg) three times per week for a total of 11 injections. (B) Racine scale scores progressively increased with repeated PTZ injections (Pearson correlation: r = 0.977, R² = 0.955, ***p < 0.001; n = 5 mice). (C) Experimental timeline after PTZ treatment (left). Electrocorticogram (ECoG) recordings were obtained 1 and 22 days after the final PTZ injection. Mice received an intravenous injection of AAV on the day following the first ECoG recording. The schematic on the right illustrates the AAV-GFP and AAV-GAD65 constructs. (D) Representative ECoG traces (a), number of seizure-like events during 15-min epochs (b; n = 5 mice; n.s., p = 0.12), and delta power (c; n = 5 mice; *p = 0.02) from PTZ-treated control mice that did not receive AAV injection. (E, F) Representative ECoG traces (a), number of seizure-like events (b; AAV-GFP, n = 4, n.s., p = 0.71; AAV-GAD65, n = 5, n.s., p = 0.08), and delta power (c; **p = 0.008 for AAV-GFP, **p = 0.007 for AAV-GAD65). Statistical comparisons were made between values obtained 1 and 22 days after the final PTZ injection within each group. (G) Group comparison of seizure-like events at 1 and 22 days; no significant differences among PTZ-treated control, AAV-GFP, and AAV-GAD65 groups. (H) Delta power at 1 and 22 days. At 22 days, AAV-GAD65 significantly reduced delta power compared with PTZ-treated control (*p = 0.01) and AAV-GFP (**p = 0.007), reaching levels comparable to naïve mice (AAV-GAD65 vs naïve, n.s., p = 0.46). (I) Scatter plots of seizure-like events versus delta power. Energy distance tests indicated a significant shift between day 1 and day 22 in the AAV-GAD65 group (**p = 0.008; n = 5 mice), with no difference compared to naïve controls (p = 0.23). Data represent mean ± SEM. Statistical significance was determined using energy distance test for equality of distributions by R.

Intravenous administration of AAV-GAD65 (Figure 4F), but not AAV-cmGAD67-GFP (AAV-GFP) (Figure 4E), two days after the final PTZ injection markedly improved cortical network activity. Both the number of seizure-like events and delta power were greatly reduced by AAV-GAD65 at 22 days (Figures 4F-b,c).

Figures 4G and 4H summarize the number of seizure-like events and delta power at 1 and 22 days after the 11th PTZ injection in the control (PTZ only), PTZ + AAV-GFP, and PTZ + AAV-GAD65 groups. In the AAV-GAD65–injected group, both parameters were markedly decreased, and delta power was significantly reduced to the level observed in naïve mice without PTZ treatment.

To further evaluate the relationship between seizure-like events and slow-wave activity, we performed a group-level analysis of seizure-like events versus delta power (Figure 4I). In PTZ-treated control mice and those receiving AAV-GFP, seizure-like events were prominent and delta power was elevated 1 days after the final PTZ injection, and both measures remained persistently abnormal at 22 days. No significant recovery was observed between the two time points, and both groups continued to differ significantly even at 22 days from naïve mice (light blue circles in Figure. 4I).

In contrast, mice treated with AAV-GAD65 exhibited a distinct trajectory. Although seizure-like events and delta power were similarly elevated 1 day after PTZ treatment, both measures showed significant recovery by 22 days. Importantly, values in the AAV-GAD65 group at 22 days were no longer significantly different from those of naïve mice (p = 0.23, Figure 4I).

These results indicate that while control and AAV-GFP groups failed to recover from PTZ-induced cortical hyperexcitability, AAV-GAD65 treatment promoted a significant normalization of both seizure-related activity and delta oscillations, restoring network activity to near-naïve levels.

To directly assess whether AAV-GAD65 enhances GABA production, we measured tissue GABA levels in the cortex and hippocampus. Biochemical assays revealed that AAV-GAD65 significantly elevated GABA content compared with control groups (Supplementary figure S8), confirming that transgene expression increased inhibitory neurotransmitter availability. These results provide mechanistic support for the observed suppression of epileptiform activity.

### AAV-GAD65 alleviates anxiety-like behavior in PTZ-kindled mice

We next investigated whether AAV-GAD65 could ameliorate anxiety-like behavior associated with PTZ kindling. In the open-field test, locomotor activity, assessed by total distance traveled, did not differ significantly among groups, indicating that AAV treatment did not alter general motor function (Figures 5A and 5B). Consistently, home-cage monitoring and rotarod/beam-walking testing confirmed that intravenous administration of AAV-GAD65 did not affect circadian activity or motor coordination (Supplementary figures S8 and S9). By contrast, the percentage of time spent in the central zone was markedly reduced in PTZ-treated control mice compared with naïve mice, reflecting increased anxiety-like behavior (Figures 5A and 5C). Importantly, this behavioral deficit was significantly reversed in mice treated with AAV-GAD65, whereas no improvement was observed in the AAV-GFP group. Notably, the central zone occupancy of AAV-GAD65–treated mice was indistinguishable from that of naïve controls.

**Figure 5.**
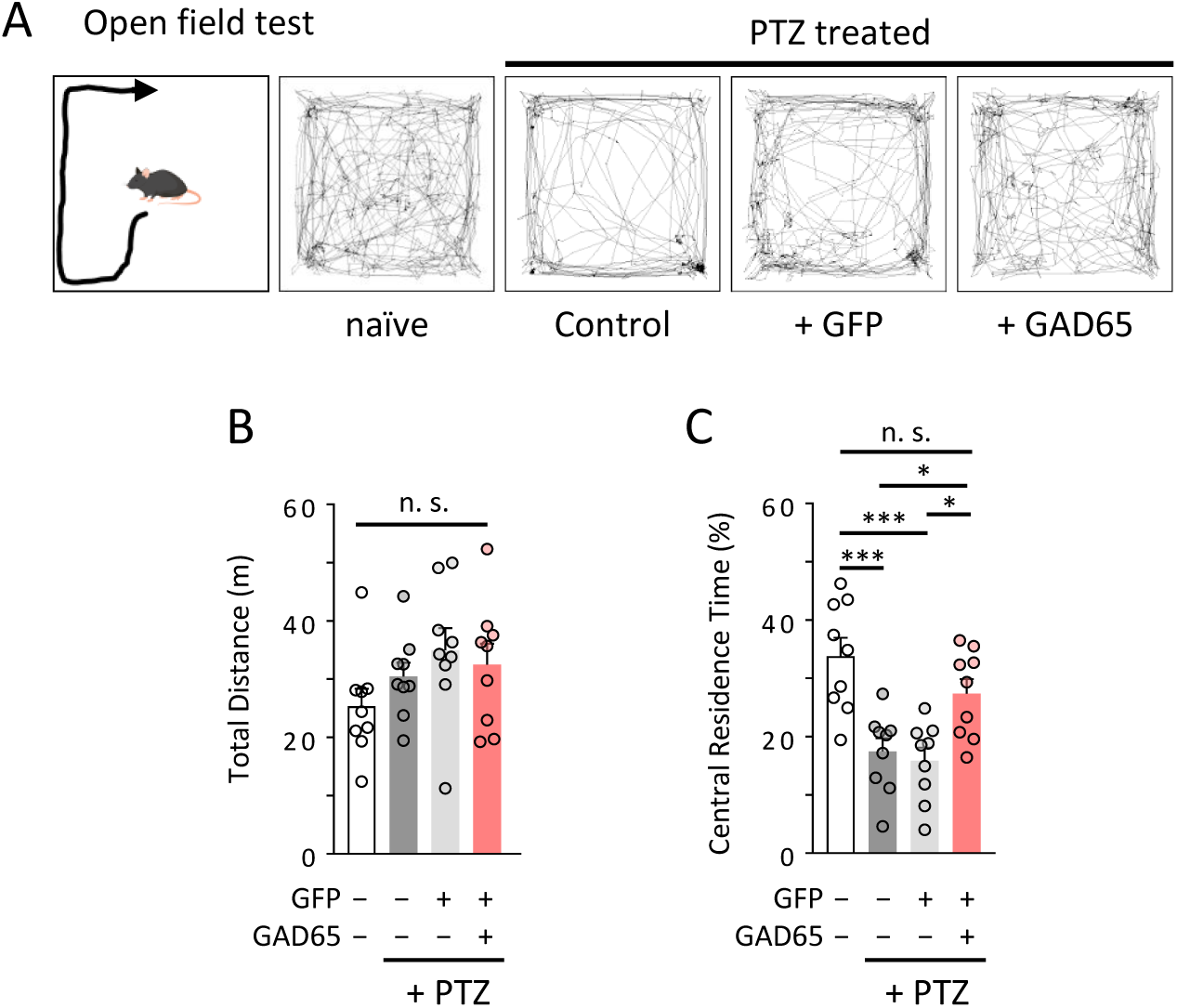
AAV-GAD65 normalizes anxiety-like behavior in PTZ-kindled mice. (A) Schematic of the open-field test (left) and representative movement trajectories of mice: naïve, PTZ-treated control, PTZ-treated +AAV-GFP, and PTZ-treated +AAV-GAD65. (B) Total distance traveled did not differ significantly among groups (n = 5 mice per group; n.s.). (C) Percentage of time spent in the central zone was significantly reduced in PTZ-treated control mice compared with naïve mice (***p < 0.001) and was restored by AAV-GAD65 treatment compared with PTZ-treated control mice (*p < 0.05). No significant difference was observed between naïve and AAV-GAD65 groups. Data represent mean ± SEM (n = 5 mice per group). Statistical significance was determined using one-way ANOVA tukey’s post hoc test.

These findings are particularly relevant in the clinical context, as anxiety is a common and debilitating comorbidity in patients with epilepsy ^14,15^. Thus, AAV-GAD65 not only suppresses epileptiform activity but also normalizes anxiety-like behavior, while maintaining normal locomotor, circadian, and motor functions, underscoring its therapeutic potential beyond seizure control.

### AAV-GAD65 attenuates seizure progression and improves survival in a severe PTZ model

To further evaluate the therapeutic efficacy of AAV-GAD65, we employed a higher-dose PTZ kindling paradigm (35 mg/kg). This paradigm is considered to mimic refractory epilepsy, as it induces more severe and persistent seizure activity than the standard 30 mg/kg protocol. In this model, Racine scale scores increased more rapidly, with significantly greater seizure severity across injection numbers compared with the 30 mg/kg group (Figures 6A and 6B). Survival analysis revealed that all mice in the 30 mg/kg group survived after 11 PTZ injections, whereas mortality in the 35 mg/kg group began at the 7th injection, resulting in 60% lethality by the end of the protocol (Figure 6C).

**Figure 6.**
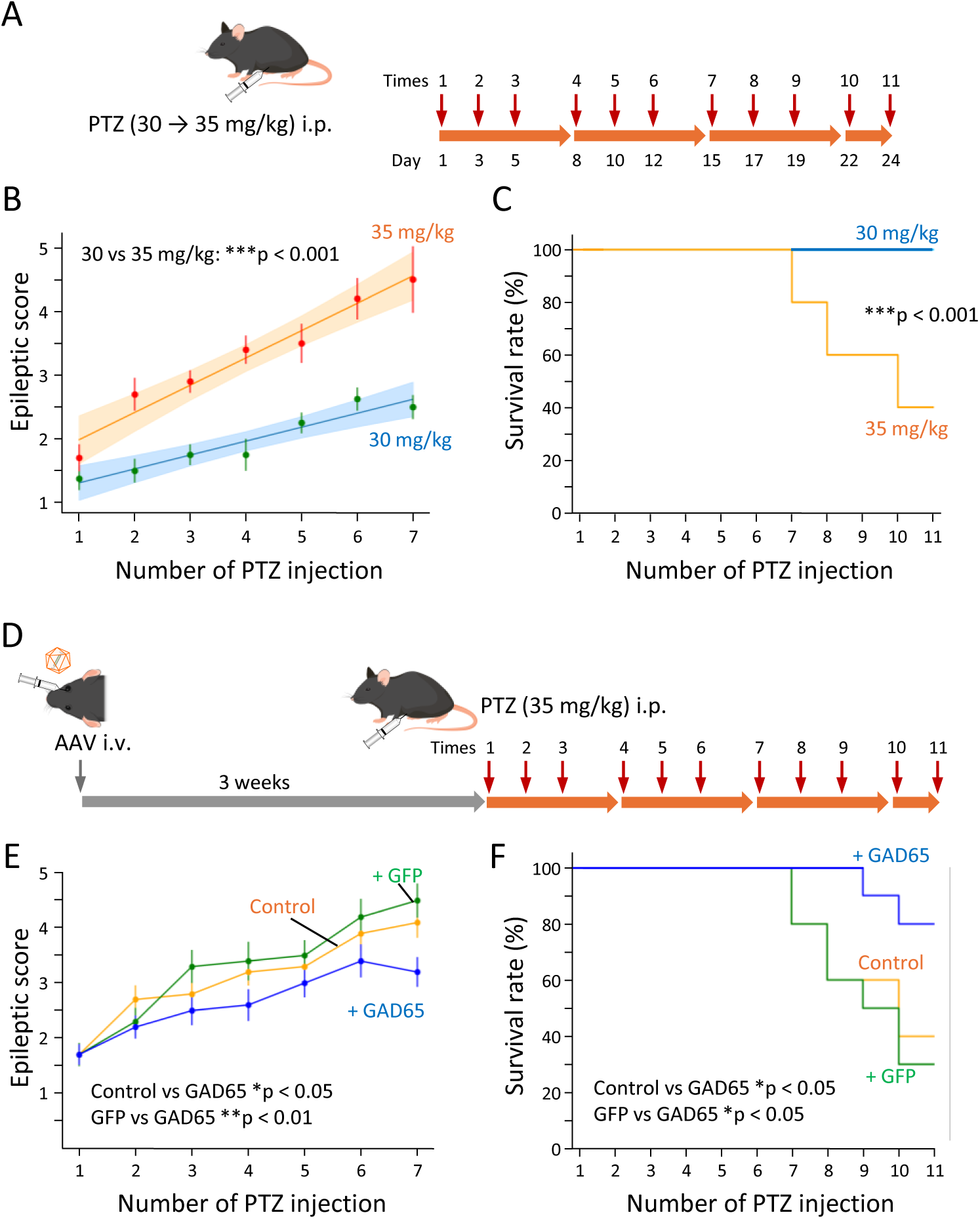
AAV-GAD65 improves seizure progression and survival in a severe PTZ model. (A) Experimental timeline for the PTZ (30 and 35 mg/kg) paradigm. (B) Racine scale scores increased in an injection number–dependent manner, with significantly higher scores in the 35 mg/kg group compared with the 30 mg/kg group (group effect, ***p < 0.001). Shaded areas represent 95% confidence intervals (CIs) of the model-derived predicted means, obtained from ordinary least squares (OLS) regression. (C) Comparison of survival curves between mice treated with PTZ at 30 mg/kg and 35 mg/kg. All eight mice survived after 11 injections at 30 mg/kg, whereas mortality began at the 7th injection in the 35 mg/kg group, and 60% of mice (6 of 10) died (***p = 0.0007, Gehan-Breslow-Wilcoxon test). (D) Schematic representation of the experimental procedure. Three weeks before the initiation of PTZ administration, AAV-PHP.eB vectors expressing either GAD65 together with GFP (AAV-GAD65) or GFP alone (AAV-GFP) were delivered intravenously. (E) In PTZ (35 mg/kg)-treated mice, seizure progression was attenuated by AAV-GAD65 (+GAD65) (*p < 0.05), whereas AAV-GFP (+GFP) had no effect (p = 0.53). (F) Survival curves showed improved survival in the AAV-GAD65 group compared with PTZ-treated controls and AAV-GFP groups (*p < 0.05, **p < 0.01, respectively) (n = 10 mice per group).

To test whether AAV-GAD65 could mitigate this severe phenotype, vectors were intravenously delivered three weeks before PTZ administration (Figure 6D). In the 35 mg/kg model, seizure progression was significantly attenuated in mice treated with AAV-GAD65, whereas AAV-GFP had no effect (Figure 6E). Moreover, survival curves demonstrated that AAV-GAD65 conferred a marked survival benefit compared with PTZ-treated controls and AAV-GFP groups (Figure 6F).

These results indicate that systemic administration of AAV-GAD65 not only reduces seizure severity but also improves survival in a stringent PTZ model, underscoring its strong therapeutic potential for refractory epilepsy.

## DISCUSSION

### Advantages of the cmGAD67 promoter for inhibitory neuron-targeted gene therapy

The development of compact, cell type–specific promoters is critical for maximizing the therapeutic potential of AAV vectors, whose limited packaging capacity restricts the inclusion of large regulatory elements. The cmGAD67 promoter, identified through a systematic analysis of the Gad1 locus, is only 410 bp in length yet drives robust transcriptional activity with high specificity for inhibitory neurons. This compact size preserves space for therapeutic payloads or additional regulatory modules, while its strength overcomes the limitations of the conventional mGAD65(delE1) promoter. The presence of the E_TSS_ core promoter and the Rii enhancer region, enriched for Dlx binding motifs, likely underlies its strong and selective activity. These features make the cmGAD67 promoter a practical platform for both experimental applications and translational gene therapy.

### Preferential targeting of PV interneurons and implications for network control

A distinguishing property of the cmGAD67 promoter is its preferential targeting of PV^+^ interneurons. PV^+^ cells mediate powerful perisomatic inhibition of pyramidal neurons and are pivotal for controlling cortical excitability ^11,12^. By biasing expression toward PV^+^ interneurons, the cmGAD67 promoter provides a unique advantage over enhancer-based strategies such as Dlx, which broadly target inhibitory neurons without subtype selectivity ^16^. Although expression in SST^+^ interneurons was also detected, off-target expression in excitatory neurons was negligible. This PV preference likely contributes to the marked efficacy of AAV-GAD65 in suppressing epileptiform activity. Notably, even if low-level expression were to occur in excitatory neurons, the enzymatic function of GAD65—converting glutamate to GABA—would be expected to reduce excitatory drive, further tipping the network toward inhibition. Thus, the promoter’s targeting profile is well suited for therapeutic strategies aimed at restoring E/I balance.

### Therapeutic effects of AAV-GAD65 in PTZ models of epilepsy

Delivery of GAD65 under the control of the cmGAD67 promoter (AAV-GAD65) produced significant therapeutic benefits in PTZ-kindled mice. Electrophysiological analyses revealed suppression of abnormal delta oscillations and reduction of seizure-like events, indicating amelioration of cortical hyperexcitability. In addition, AAV-GAD65 normalized anxiety-like behavior in the open-field test, an outcome of clinical relevance given the high prevalence of psychiatric comorbidities in epilepsy ^14,15^. Importantly, biochemical assays confirmed that AAV-GAD65 elevated GABA levels in both cortex and hippocampus (Supplementary figure S8), directly linking its therapeutic effects to enhanced inhibitory neurotransmitter synthesis. Furthermore, in a severe PTZ paradigm mimicking treatment-resistant epilepsy, AAV-GAD65 not only attenuated seizure progression but also conferred a clear survival advantage.

It should be noted that the 35 mg/kg PTZ paradigm does not fully recapitulate pharmacoresistant epilepsy, clinically defined as the failure of multiple antiseizure drugs. Rather, it provides a model of severe, recurrent seizures with high mortality, thereby mimicking key features of treatment-resistant epilepsy. This distinction is important when interpreting the translational relevance of our findings.

### Translational potential and limitations of the present approach

The cmGAD67 promoter offers a strong foundation for inhibitory neuron-targeted gene therapy. Its compactness and high specificity make it compatible with systemic AAV delivery, particularly in combination with blood–brain barrier-penetrant capsids. Nevertheless, several limitations should be acknowledged. First, the current study was limited to PTZ-induced epilepsy models; validation in other chronic epilepsy models and in larger brains will be essential to establish broader translational relevance. Second, while off-target expression was negligible in mice, thorough evaluation of safety and cell-type specificity will be required in nonhuman primates and ultimately in humans. Third, peripheral leakage of systemically delivered AAVs was not assessed here and represents a potential safety concern for clinical translation.

Although the PTZ paradigm mimics several aspects of treatment-resistant epilepsy, further validation in chronic and drug-resistant models will be required to strengthen translational claims. Importantly, systemic administration of AAV-GAD65 did not disrupt circadian activity or motor coordination (Supplementary figures S9 and S10), supporting a favorable safety profile in addition to its therapeutic efficacy. The translational feasibility of GAD-based therapy is further supported by a first-in-human clinical trial in Parkinson’s disease, where AAV-GAD65 delivery to the subthalamic nucleus demonstrated both efficacy and safety ^10^. Together, these findings highlight AAV-cmGAD67-GAD65 as a promising gene therapy platform for treatment-resistant epilepsy and potentially other disorders characterized by E/I imbalance.

### Broader implications for disorders involving E/I imbalance

Beyond epilepsy, disruption of the E/I balance is implicated in a wide spectrum of brain disorders. Reduced inhibitory function has been linked to anxiety disorders, schizophrenia, and autism spectrum disorders, all of which are associated with network hyperexcitability ^1–3^. Moreover, accumulating evidence suggests that impaired GABAergic transmission contributes to the progression of neurodegenerative diseases, including Alzheimer’s disease ^17–21^, where loss of inhibitory tone exacerbates aberrant oscillatory activity and cognitive decline. The cmGAD67 promoter, by enabling compact and selective gene delivery to inhibitory neurons—particularly PV^+^ interneurons—provides a versatile tool to explore circuit mechanisms and to develop therapeutic interventions. Thus, AAV vectors equipped with the cmGAD67 promoter may serve not only as a disease-modifying approach for treatment-resistant epilepsy but also as a broadly applicable platform for correcting circuit dysfunction across diverse neurological and psychiatric conditions characterized by E/I imbalance.

## CONCLUSION

We developed a compact 410-bp GAD67 promoter (cmGAD67) that drives strong and selective expression in inhibitory neurons, with preferential targeting of PV^+^ interneurons. Systemic AAV delivery of cmGAD67-GAD65 effectively suppressed epileptiform activity, normalized anxiety-like behavior, and improved survival in PTZ models. These findings establish the cmGAD67 promoter as a versatile platform for inhibitory neuron-targeted gene therapy, with strong therapeutic potential for treatment-resistant epilepsy and broader applicability to disorders involving E/I imbalance.

## MATERIALS & METHODS

### Animals

All animal experiments were approved by the Animal Care and Use Committee of Gunma University (Approval Nos. 24–053 and 24–057) and conducted in accordance with the Japanese Act on the Welfare and Management of Animals and the Guidelines for Proper Conduct of Animal Experiments issued by the Science Council of Japan. Wild-type C57BL/6J mice and VGAT-tdTomato mice (Riken BRC no. RBRC10799) were used for experiments. Mice of both sexes, aged 6–10 weeks, were housed under a 12-h light/dark cycle with ad libitum access to food and water. Care was taken to avoid sex-related bias, and all efforts were made to minimize animal suffering and reduce the number of animals used.

### AAV Vector Construction and Production

Recombinant AAV vectors were generated by cloning transgene cassettes into AAV2 ITR-containing plasmid backbones. Inhibitory neuron-specific promoters used in this study included the conventional mGAD65 promoter (2,542 bp), the compact mGAD65 (cmGAD65, 486 bp), and the compact mGAD67 (cmGAD67, 410 bp), each driving the expression of reporter (GFP, tdTomato) or effector genes (ChR2(H134R)-EGFP ^22^, hM4D(Gi) (Addgene plasmid #50461), or mGAD65). For cell-type-selective dual expression, a 0.4 kb mCaMKII promoter ^23^ was used for excitatory neuron targeting.

AAV vectors were produced in HEK293T cells by triple transfection of the expression plasmid (pAAV), adenoviral helper plasmid (pHelper, Agilent Technologies), and a Rep/Cap plasmid using polyethylenimine Max (Polysciences) as described previously ^24^. Briefly, cells were cultured in DMEM supplemented with 8% fetal bovine serum. Six days after transfection, the culture medium was harvested, and viral particles were precipitated using 8% polyethylene glycol 8000 and 500 mM NaCl, followed by resuspension in D-PBS. Viral preparations were purified by iodixanol (Optiprep; AXS-1114542-250ML, Alere Technologies AS, Oslo, Norway) density gradient ultracentrifugation and concentrated using Vivaspin Turbo 15 MWCO 100000 PES (VS15T42; Sartorius, Göttingen, Germany). Final titers were measured by quantitative PCR using Power SYBR Green PCR Master Mix (Thermo Fisher Scientific) with primers targeting the WPRE sequence: forward, 511-CTGTTGGGCACTGACAATTC-311; reverse, 511-GAAGGGACGTAGCAGAAGGA-311.

### Intravenous and Stereotactic AAV Injection

#### Intravenous Injection

Mice were anesthetized with intraperitoneal ketamine (100 mg/kg body weight) and xylazine (10 mg/kg). Anesthetic depth was verified via the toe-pinch reflex. A total of 100 μL of AAV solution was slowly injected into the retro-orbital venous sinus using a 30-gauge needle (Nipro Co., Osaka, Japan).

#### Stereotactic Injection

Mice were anesthetized with intraperitoneal ketamine (70 mg/kg body weight [BW]) and xylazine (7.0 mg/kg BW) and maintained under 0.7% isoflurane anesthesia using an anesthetic vaporizer (MK-AT210; Muromachi Kikai, Fukuoka, Japan). Anesthetic depth was monitored via the toe-pinch reflex. A hole was created over the injection site using a 30G needle. AAV vectors were bilaterally injected into the hippocampus. A 10-μL Hamilton syringe with a 33G needle was used with a stereotaxic micromanipulator (SMM-100; Narishige, Tokyo, Japan) mounted on a stereotactic frame (SRS-5-HT; Narishige). The stereotaxic coordinates relative to bregma; AP −2.0 mm, ML ±0.75mm, DV +1.7 mm (advanced to +1.9 mm, then retracted by −0.2 mm).

### Brain Tissue Preparation and Imaging

At 3 weeks post-injection, mice were transcardially perfused with PBS followed by 4% paraformaldehyde (PFA). Brains were post-fixed for 6 hours and coronally sectioned at 50 μm thickness using a vibratome (VT1200S; Leica, Wetzlar, Germany). For electrophysiology, acute slices (250–300 μm) were prepared. Fluorescence images were acquired using a fluorescence microscope (VB-7010 or BZ-X800; Keyence, Osaka, Japan) or a confocal microscope (LSM 800; Zeiss, Oberkochen, Germany). Image analysis was performed using ZEISS ZEN and/or Fiji (https://fiji.sc/).

### Quantification of GFP Fluorescence and Cell Type Specificity

Whole-brain GFP fluorescence was quantified using Fiji software. Fluorescence intensity was quantified by measuring the brightness of ROIs in cerebral cortex excluding the olfactory bulb, superior colliculus, and cerebellum. To assess cell-type specificity, GFP⁺ cells co-labeled with tdTomato (VGAT-tdTomato) or with marker antibodies (e.g., parvalbumin [PV], somatostatin [SST]) were manually counted using a BZ-X800 widefield microscope (Keyence). Co-localization rates were calculated for each section and then averaged per animal. Cell counting was performed in the M1–M2 region of the cerebral cortex.

### Immunohistochemistry

Free-floating brain sections were incubated for 30 min at Room Temperature (RT) in a blocking solution containing; 5% normal donkey serum, 0.5% Triton X-100, and 0.05% sodium azide in phosphate-buffered saline (PBS). Primary antibodies in blocking solution were applied and incubated overnight at 4 °C with gentle shaking.

For Figure 2, the following primary antibodies were used:

- Mouse anti-GFP (1:1,000; GTX21218, GeneTex, CA, USA)
- Goat anti-parvalbumin (1:200; ab11427, Abcam, Cambridge, UK)
- Rat anti-somatostatin (1:100; MAB354, Millipore, MA, USA)

After two washes with 0.5% Triton and three washes with 0.1% Triton, sections were incubated for 3h at RT with gentle shaking with fluorophore-conjugated secondary antibodies diluted in the same blocking solution:

- Alexa Fluor 488-conjugated donkey anti-mouse IgG (1:2,000; A32766, Invitrogen, MA, USA)
- Alexa Fluor 555-conjugated donkey anti-goat IgG (1:2,000; A32816, Invitrogen)
- Alexa Fluor 647-conjugated donkey anti-rat IgG (1:2,000; A448272, Invitrogen)

After two washes with 0.5% Triton, three washes with 0.1% Triton and two washes with PBS, sections were mounted using ProLong Diamond (Thermo Fisher Scientific, MA, USA). Co-localization of GFP with these markers was analyzed by BZ-800 and confocal microscopy.

For Figures 3E and 3F, the following primary and secondary antibodies were used:

- Rat anti-GFP (1:1,000; 04404-84, Nacalai Tesque, Kyoto, Japan)
- Rabbit anti-GABA (1:250; ABN131, Millipore)
- Alexa Fluor 488-conjugated donkey anti-rat IgG (1:2,000; A48269, Invitrogen)
- Alexa Fluor 555-conjugated donkey anti-rabbit IgG (1:2,000; A32794, Invitrogen)

### Electrophysiology

Patch-clamp recordings were performed in acute cerebral slices of the mouse motor cortex including primary and secondary motor areas, as described previously ^25,26^ with some modifications. Three to five weeks after the intravenous injection of a BBB-permeable AAV vector expressing ChR2(H134R)-EGFP-P2A-NLS-EGFP ^22^ under the control of cmGAD67 promoter into the VGAT-tdTomato mice as described elsewhere, coronal slices (250-300 µm in thickness) of the cerebral cortex containing the motor areas were prepared using a vibroslicer (VT1200S; Leica). After a two-step incubation of the slices, electrophysiological recordings were conducted ^25^. The extracellular solution (artificial cerebrospinal fluid) contained (in mM): 125 NaCl, 2.5 KCl, 2 CaCl_2_, 1 MgCl_2_, 1.25 NaH_2_PO_4_, 26 NaHCO_3_, and 10 D-glucose, and was continuously bubbled with 95% O_2_ and 5% CO_2_ gas mixture. Confocal fluorescence signals of the acute brain slices were acquired using a water-cooled EM-CCD camera (512 × 512 pixels; iXon3 DU-897E-CS0-#BV-500; Andor, Belfast, Northern Ireland), a 40× water immersion objective (LUMPLFLN 40XW; Olympus, Tokyo, Japan) and a high-speed spinning-disk confocal unit (CSU-X1; Yokogawa Electric, Tokyo, Japan) mounted to an upright microscope (BX51WI; Olympus, Tokyo, Japan). A 488 nm blue light beam from a diode laser module (Stradus 488-50, VORTRAN, USA) and a 561 nm laser (LDSYS-561GH-SP43, Solution Systems, Japan) were used as excitation light for GFP and tdTomato, respectively. Emitted fluorescence was collected through a band-pass filter (500-550 nm for GFP and 580-660 nm for tdTomato). For ChR2 activation, the 488 nm blue light used for GFP excitation was applied to the slice for ∼400 ms via the confocal unit.

Whole-cell recordings were performed at room temperature from inhibitory neurons (i.e. tdTomato-positive cells) in the motor cortex using patch pipettes (1–4 M) pulled from borosilicate glass (Harvard Apparatus) or #0010 glass (PG10165-4, World Precision Instruments). ChR2-mediated inward photocurrents were recorded in voltage-clamp mode at a holding potential of −60 mV. Passive membrane properties of the recorded cells were assessed by applying voltage pulses from −60 to −65 mV for 500 ms. To record light-induced firing of action potentials, current clamp recordings were performed, and the resting potential was adjusted to −60 mV by current injection. In some cases, cell-attached extracellular recordings (or loose patch recordings) were conducted prior to establishing whole-cell recordings (i.e. before rupturing the patch membrane), to examine ChR2-mediated light-induced spike discharges. All the electrical signals were low pass filtered at 10 kHz and sampled at 50 kHz with Digidata 1440A (Molecular Devices) and the pCLAMP10 software. The liquid junction potentials were not corrected. Data analysis and statistical tests were performed using Igor Pro9 (Wavemetrics) with Neuromatic (http://www.neuromatic.thinkrandom.com) ^27^ and GraphPad Prism10 software.

### ECoG Signal Analysis

For DREADD experiments (Figures 3E–3G), continuous ECoG was recorded for 30 min immediately before and 30 min after DCZ administration. ECoG was recorded in mice three weeks after AAV vector injection, both before and after intraperitoneal administration of 1.0 mg/kg DCZ (Cosmo Bio, Tokyo, Japan) as previously described ^28^. For PTZ experiments (Figure. 4), 30 min recordings were acquired on day 1 and 22.

Screw electrodes were placed on the cortical surface with the following coordinates: recording electrode—AP: −2.00 mm, ML: +2.00 mm from Bregma; reference electrode—AP: −1.50 mm, ML: +1.50 mm from Lambda; ground electrode—AP: −1.50 mm, ML: −1.50 mm from Lambda. Electrodes were secured and insulated using GC Unifast III (GC Corporation, Tokyo, Japan). ECoG signals were amplified x 2,000 using a HAS-4 headstage amplifier (ADInstruments, Dunedin, New Zealand), then digitized using a PowerLab data acquisition system (ADInstruments). The signals were recorded at a sampling rate of 1 kHz and high-pass filter at 0.5 Hz and a low-pass filter at 60 Hz.

ECoG analyses used the final 15 min of each 30-min recording. Spike counts (Number / 15 min) were obtained by detecting waveforms with absolute peak amplitude between 300–1000 μV. Delta power was computed as the mean power within 0.5–4.0 Hz (μV²). ECoG data were acquired and visualized using LabChart software (ADInstruments), and waveforms were further analyzed with MATLAB R2024b (MathWorks, MA, USA).

### Open field test

The TimeOFCR1E system (O’Hara & Co., Ltd., Tokyo, Japan) was used. Mice were released in the center of a 40 cm × 40 cm field under 40 lux illumination, and their behavior was observed for 10 minutes. Prior to testing, mice were handled for 5 minutes per day for 1 week to habituate them to the experimenter.

### PTZ injection and seizure induction

Seizures were induced with pentylenetetrazole (PTZ; Cayman Chemical, Michigan, USA) based on previous reports ^29,30^. PTZ was freshly prepared on the day of each experiment and administered intraperitoneally to 7-week wild-type mice (30–35 mg/kg) 3×/week for 11 sessions.

### Epilepsy score measurement

Following each PTZ injection, seizure behavior was monitored for 20 minutes. Seizure severity was assessed using a 7-point epilepsy scoring scale (0–6), based on established criteria ^29,30^. Survival was monitored daily.

### Statistical analysis

Data were analyzed using GraphPad Prism (version 7 or 10) or R (version 4.5.1). Results are presented as mean ± standard error of the mean (SEM).

## RESOOURCE AVAILABILITY

### LEAD CONTACT

Further information and requests for resources and reagents should be directed to and will be fulfilled by the lead contact, Hirokazu Hirai.

### MATERIALS AVAILABILITY

The cmGAD67 promoter described in this study is the subject of patent applications. Plasmids of the cmGAD67 promoter have been deposited to Addgene and is available to the research community.

### DATA AVAILABILITY

All data supporting the findings of this study are available within the article and its supplementary files. Custom MATLAB scripts used for EEG spectral analysis are available from the lead contact upon reasonable request.

## ACKNOWLEDGMENTS

This work was supported by grants from the Program for Brain Mapping by Integrated Neurotechnologies for Disease Studies (Brain/MINDS; JP20dm0207057/JP21dm0207111 [to H.H.]) and Multidisciplinary Frontier Brain and Neuroscience Discoveries (Brain/MINDS 2.0; JP24wm0625103 [to H.H.]) from the Japan Agency for Medical Research and Development (AMED); and by MEXT/JSPS KAKENHI (20K06906/24K10022 [to N.H.], 22K06454/24H01221 [to A.K.], and 23H02791 [to H.H.]) and Next-GIP (JPMJSP2146 [to Y.F.]). The authors thank Dr. Shigeo Miyata for the advice on PTZ-induced epilepsy models; Asako Ohnishi, Nobue McCullough, Chieko Miyazawa, and Ayako Sugimoto and Keiko Sato for AAV vector production; Junko Sugiyama for mouse care; Hiroyoshi Inaba for assistance with EEG recordings; and Mamiko Suzuki for assistance with open field testing.

## AUTHOR CONTRIBUTIONS

Y.F., A.K., N.H., and H.H. designed the experiments. Y.F. performed most of the experiments. N.H. conducted the slice patch-clamp recordings. K.M. performed direct AAV injections into the mouse hippocampus. R.K. provided the VGAT-tdTomato mice. A.K. supervised the production of various AAV vectors. H.H. supervised and finalized the entire study.

## SUPPLEMENTAL INFORMATION

Supplementary figures S1–S10

Video S1. DCZ-induced seizure in a mouse with AAV-driven Gi-DREADD expression in hippocampal inhibitory neurons

