## Supplementary Information for "A compact glutamic acid decarboxylase 67 promoter enables inhibitory neuron-targeted AAV gene therapy for treatment-resistant epilepsy"

### Supplementary figure 1

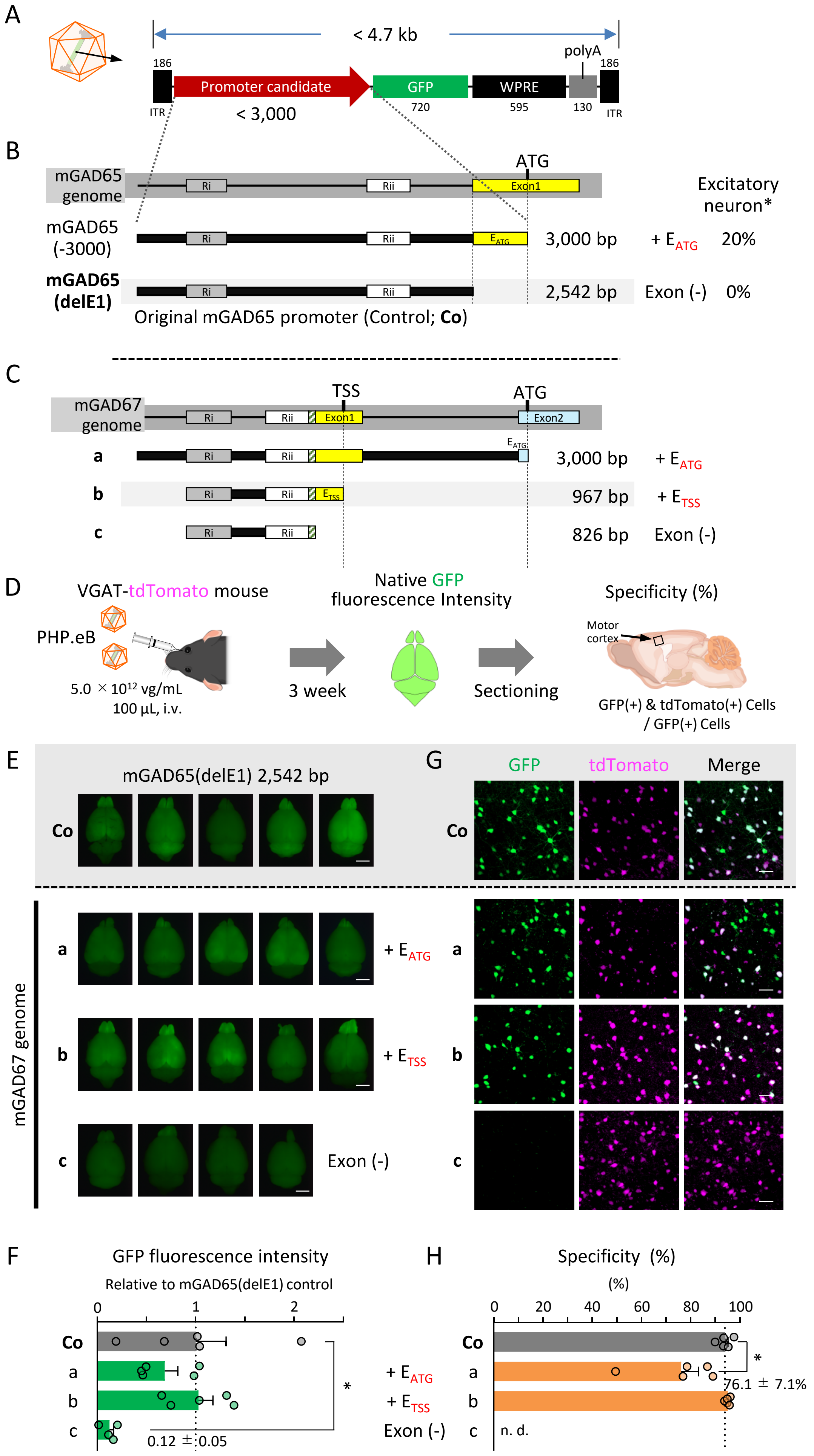

**Supplementary figure 1. Strategy for identifying a minimal promoter region specific to GABAergic neurons.**

A. Schematic of the AAV genome construct used for screening candidate GABAergic neuron-specific promoters.

B. Schematic representation of the mGAD65 genomic fragments used in a previous study for identifying GABAergic neuron-specific promoters [1]. In that study, when a 3-kb upstream fragment from the first ATG of the mGAD65 (*Gad2*) gene (referred to as mGAD65(-3000)) was used to drive GFP expression in AAV-PHP.B and systemically delivered to mice, approximately 20% of GFP-positive cells were identified as excitatory neurons. In contrast, deletion of the Exon 1 region up to the ATG (EATG) (mGAD65(delE1), 2542 bp) eliminated GFP expression in excitatory neurons, resulting in nearly 100% specificity for GABAergic neurons.

C. Schematic representation of candidate GABAergic neuron-specific promoter constructs cloned from the mGAD67 (*Gad1*) genomic locus. A 3-kb upstream fragment from the first ATG located in Exon 2 was selected as the initial candidate promoter (a), followed by stepwise truncation. A 967-bp fragment (b) spanning from the 5′ end of Ri to the transcription start site (TSS) within Exon 1 was generated. Additionally, an 826-bp fragment (c) lacking the $E_{TSS}$ (the Exon 1 segment up to the TSS) was prepared.

**D**. Schematic of the experimental procedure. AAV-PHP.eB vectors expressing GFP under either one of the promoter candidates (depicted in C) were intravenously injected into VGAT-tdTomato mice. Three weeks later, animals were perfusion-fixed and whole-brain GFP fluorescence was measured. Sagittal brain sections were then prepared, and GFP expression specificity in inhibitory neurons of the motor cortex was evaluated.

**E**. Whole-brain GFP fluorescence images from mice intravenously injected with PHP.eB vectors expressing GFP under the control of the control mGAD65(delE1) promoter (Co) or mGAD67 promoter candidates (a–c). Scale bars, 5 mm.

**F**. Quantification of whole-brain GFP fluorescence intensity. Fluorescence intensity in the control mGAD65(delE1) group (Co) was normalized to 1. The absence of the $E_{TSS}$ (c) led to a marked reduction in GFP expression.

**G**. Representative native fluorescence images of GFP and tdTomato in the motor cortex, along with merged images. Scale bars, 50 µm.

**H**. Quantification of GFP expression specificity in GABAergic neurons of the motor cortex. Each dot in the graphs in graphs **C** and **E** represents data from an individual mouse. Inclusion of the $E_{ATG}$ region reduced GABAergic neuron specificity (**a**), whereas constructs ending at the $E_{TSS}$ (**b**) did not. *$p < 0.05$ by one-way ANOVA with Dunnett’s post hoc test. n.d., not determined.

### Supplementary figure 2

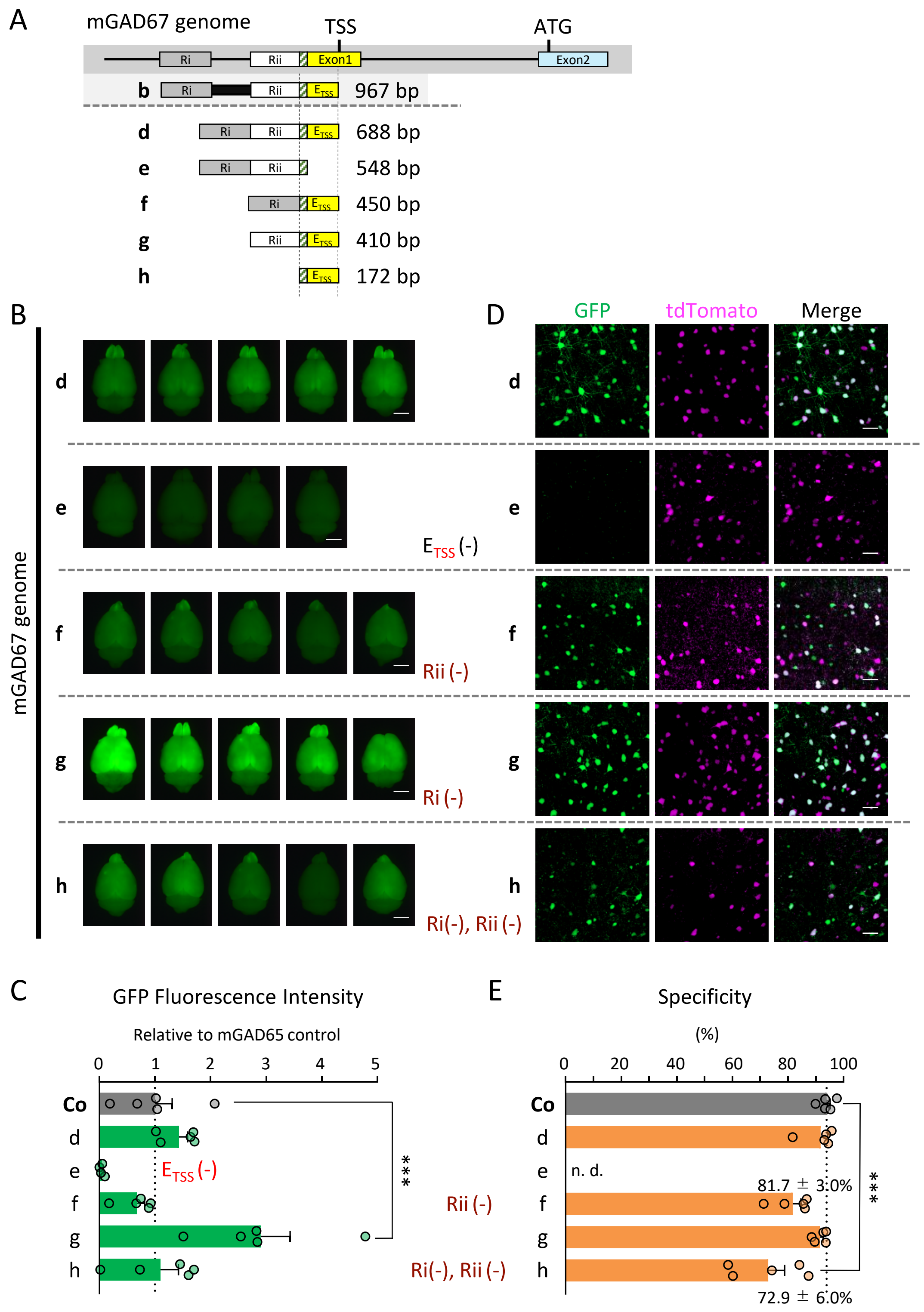

**Supplementary figure 2. The Rii enhancer and $E_{TSS}$ from the GAD67 genomic locus are sufficient to drive GABAergic neuron-specific expression.**

**A**. Schematic of the mGAD67 (*Gad1*) genomic region and the various deleted promoter candidates. Ri and Rii indicate enhancer elements reported to contain multiple binding sites for GABAergic neuron-specific Dlx transcription factor.

**B**. Whole-brain GFP fluorescence images from VGAT-tdTomato mice intravenously injected with AAV-PHP.eB vectors expressing GFP under the control of a genomic fragment containing Ri–Rii–$E_{TSS}$ (**d**), or variants with individual elements deleted (**e**–**h**), analyzed 3 weeks post-injection. Scale bars, 5 mm.

**C**. Quantification of whole-brain GFP fluorescence intensity corresponding to the conditions in panel **B**. GFP fluorescence driven by the mGAD65(delE1) promoter (Co) was normalized to 1.

**D**. Native fluorescence images of GFP and tdTomato in sagittal brain sections of the motor cortex. Green: GFP; magenta: tdTomato (inhibitory neurons). Scale bars, 50 µm.

**E**. Quantification of the proportion of GFP-positive cells that were also tdTomato-positive (indicative of GABAergic neuron specificity). Each dot represents data from an individual mouse. ***$p < 0.001$ by one-way ANOVA with Dunnett’s post hoc test. n.d., not determined.

### Supplementary figure 3

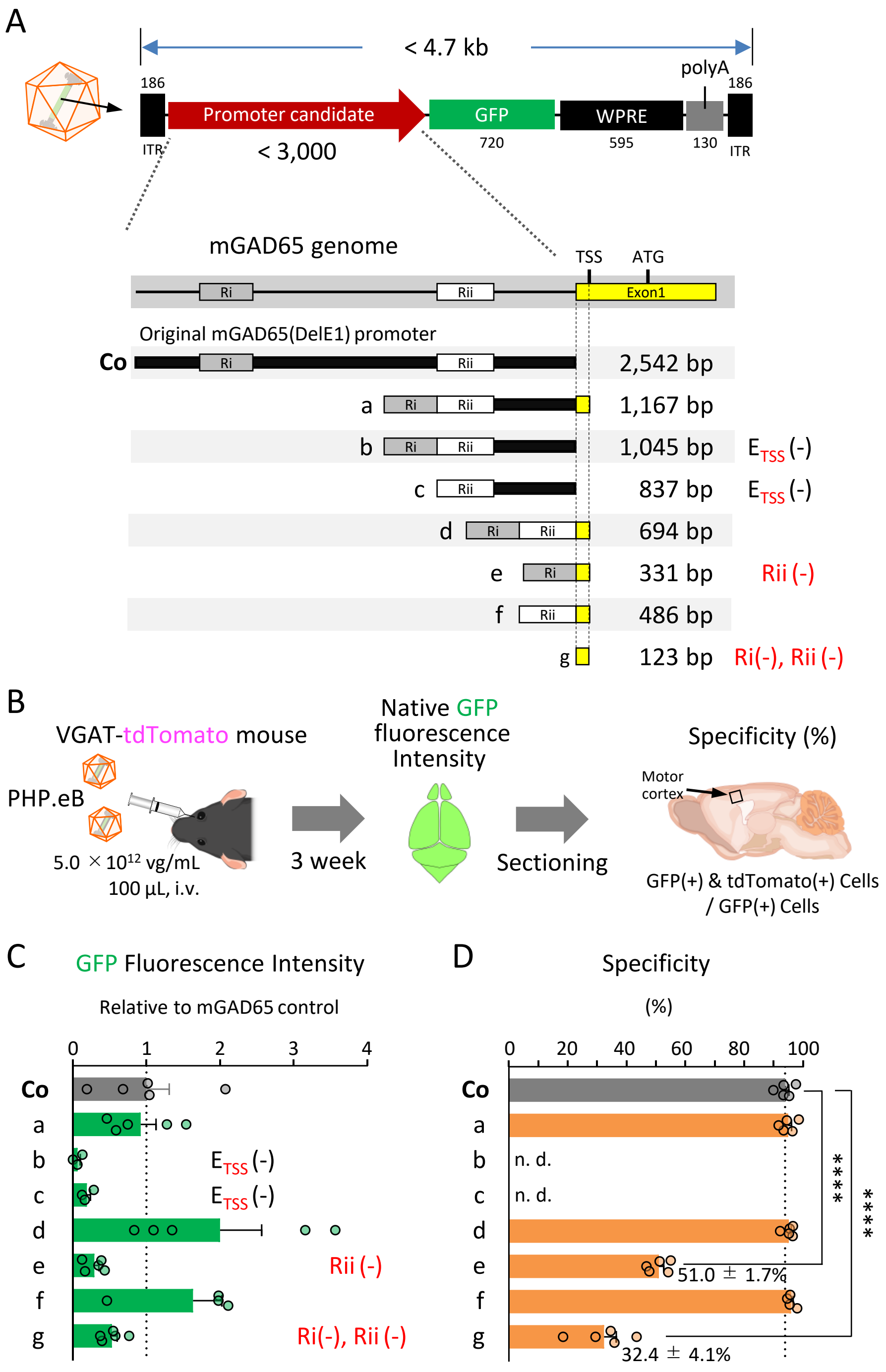

**Supplementary figure 3. A hybrid promoter composed of Rii and $E_{TSS}$ from the mGAD65 locus also drives GABAergic neuron-specific expression.**

**A**. Schematic representation of the control mGAD65(delE1) promoter (Co) and its deletion constructs (a–g). Various constructs were truncated while retaining all or some of the Ri, Rii, and $E_{TSS}$ elements.

**B**. Schematic of the experimental procedure. AAV-PHP.eB vectors expressing GFP under the control of mGAD65(delE1) or its deletion constructs were intravenously injected into VGAT-tdTomato mice. Three weeks later, whole-brain GFP fluorescence was measured, followed by preparation of sagittal brain sections to evaluate GABAergic neuron specificity in the motor cortex.

**C**. Quantification of whole-brain GFP fluorescence intensity. The fluorescence intensity obtained with the mGAD65(delE1) control promoter was normalized to 1.

**D**. Quantification of GABAergic neuron specificity. The proportion of GFP-positive cells that were also tdTomato-positive (i.e., GABAergic neurons) was calculated in the motor cortex. Each dot represents data from an individual mouse. Constructs containing both Rii and $E_{TSS}$ drove GFP expression at levels comparable to or higher than the control, while maintaining high specificity for inhibitory neurons. ****$p < 0.0001$ by one-way ANOVA with Dunnett's post hoc test. n.d., not determined.

Supplementary figure 4

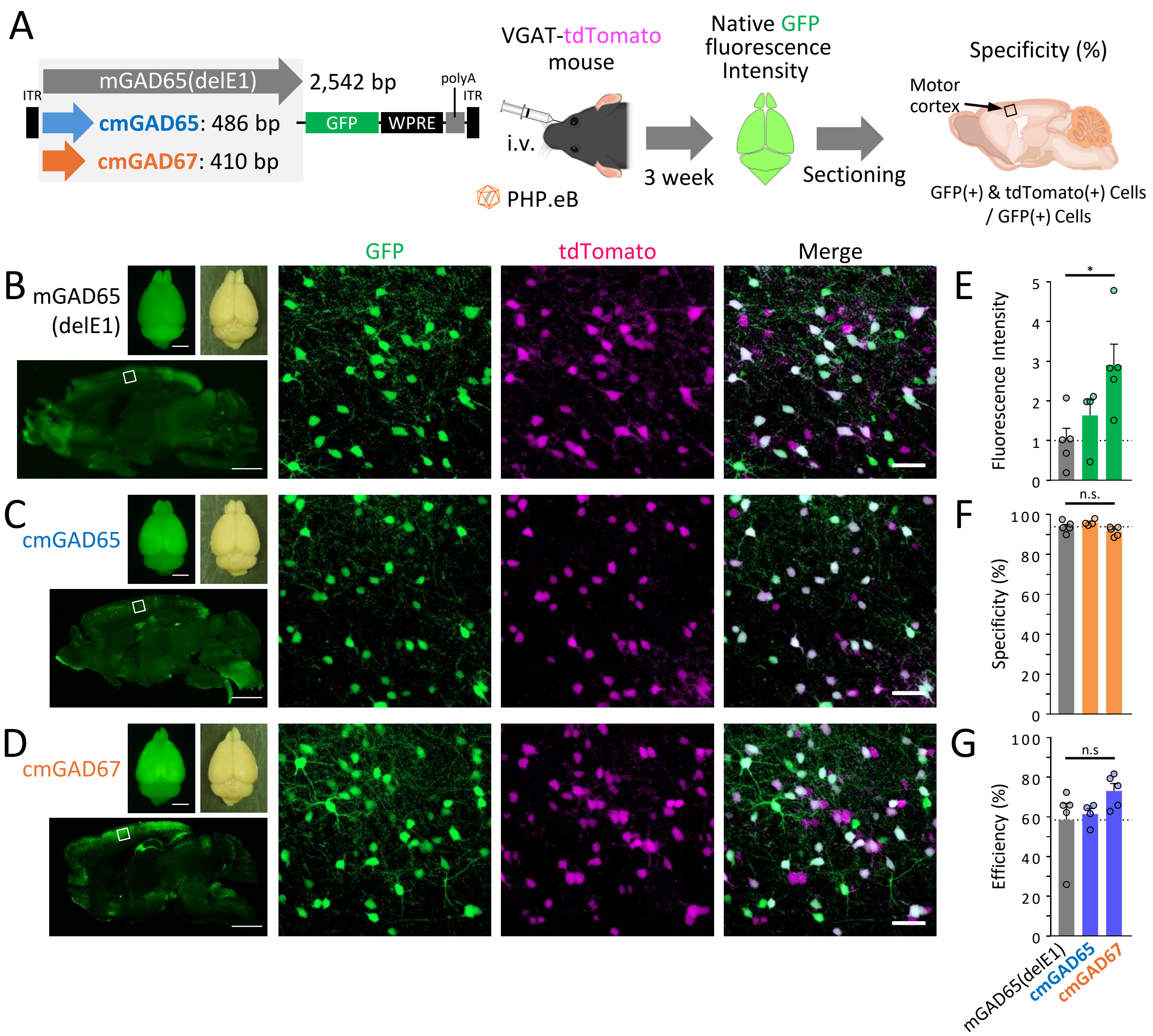


**Supplementary figure 4. Comparative analysis of expression profile of mGAD(delE1) and cmGAD promoters after intravenous administration of AAV-PHP.eB**

**A.** Schematic representation of the experimental procedure. AAV-PHP.eB expressing GFP under the control of the conventional mGAD65(delE1) promoter, cmGAD65 promoter, or cmGAD67 promoter was intravenously administered to *VGAT*-tdTomato mice. Three weeks post-injection, whole-brain GFP fluorescence intensity was measured. Sagittal brain sections were then prepared to assess the specificity of GFP expression in inhibitory neurons within the primary motor cortex.

**B-D.** Representative images of GFP fluorescence and bright-field views of the whole brain (top left paired images) and sagittal section GFP fluorescence images (bottom left) for the mGAD65(delE1) (**B**), cmGAD65 (**C**), and cmGAD67 (**D**) promoters. The three rightmost columns show magnified fluorescence images of the primary motor cortex (corresponding to the boxed region in the bottom left images). Scale bars, 5 mm (whole brains), 200 µm (sagittal sections) and 50 µm (right panels).

**E.** Quantification of whole-brain GFP fluorescence intensity except for the cerebellar area. Relative intensities are shown with the mGAD65(delE1) promoter set as 1.

**F.** Graph showing the specificity of GFP expression in inhibitory neurons.

**G.** Quantification of the proportion (efficiency) of inhibitory neurons in the primary motor cortex expressing GFP. Dots in the graphs represent individual mouse data points. *$p < 0.05$, n.s., not significant by one-way ANOVA with Dunnett's post hoc test.

Supplementary figure 5

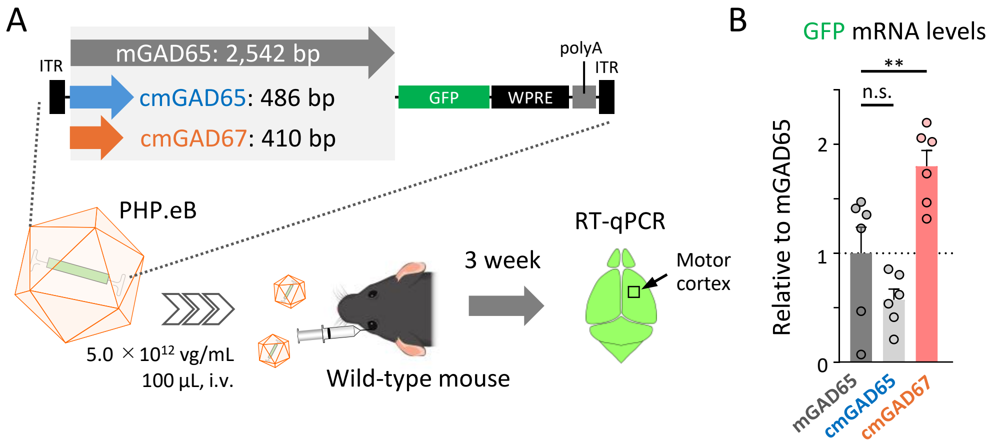


**Supplementary figure 5. The cmGAD67 promoter exhibits significantly higher transcriptional activity than mGAD65(delE1).**

A. Schematic of the experimental procedure. AAV-PHP.eB vectors ($5.0 \times 10^{12}$ vg/mL, 100 μL) expressing GFP under the control of mGAD65(delE1), the cmGAD65 promoter, or the cmGAD67 promoter were intravenously injected into VGAT-tdTomato mice. Three weeks later, total RNA was extracted from the motor cortex, and GFP mRNA levels were quantified using RT-qPCR.

B. Quantification of GFP mRNA levels. Values are shown relative to those obtained with the mGAD65(delE1) promoter, which was normalized to 1. Each dot represents data from an individual mouse. $^{**}p < 0.01$, n.s.; not significant by one-way ANOVA with Dunnett's post hoc test.

Supplementary figure 6

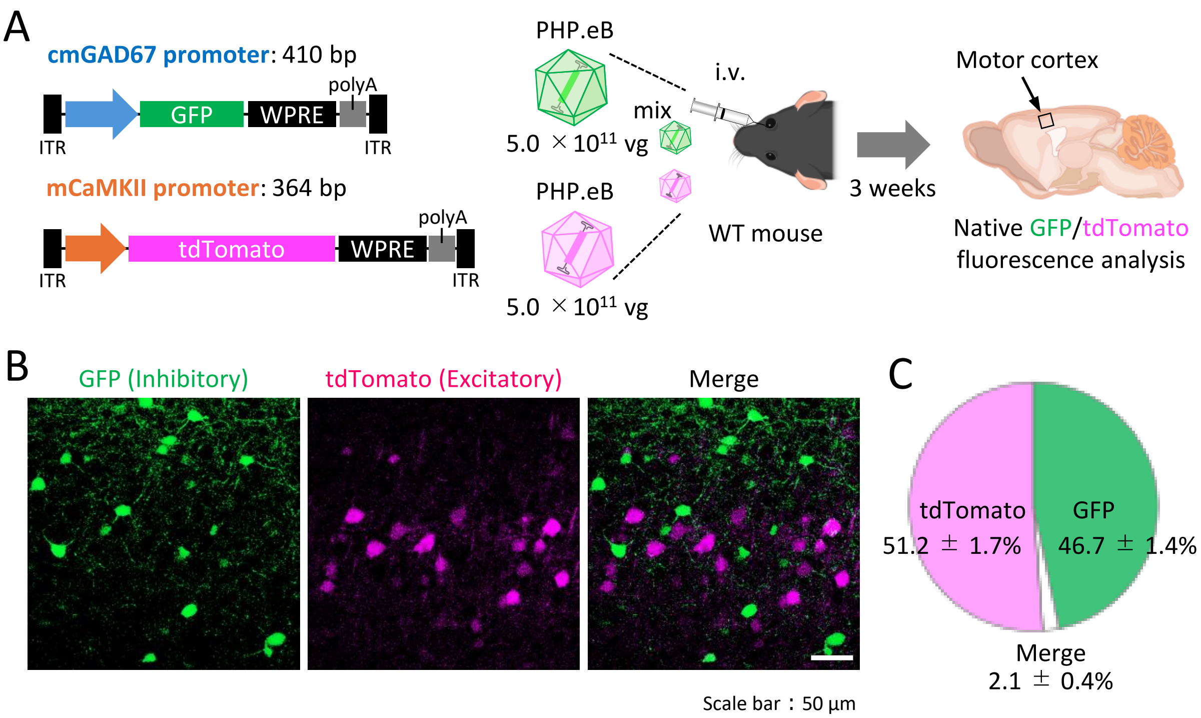


**Supplementary figure 6. Validation of inhibitory vs excitatory neuron targeting using co-injection of cmGAD67-GFP and CaMKII-tdTomato AAVs.**

**A.** Schematic representation of the experimental procedure. AAV-PHP.eB expressing GFP under the control of the cmGAD67 promoter and AAV-PHP.eB expressing tdTomato under the control of the mCaMKII promoter were mixed in equal volumes (50 µL each, $1.0 \times 10^{13}$ vg/mL) and administered via the orbital venous sinus. Three weeks post-injection, sagittal brain sections were prepared, and fluorescence microscopy was used to observe the primary motor cortex.

**B.** Fluorescence images of the motor cortex. Scale bar, 50 µm.

**C.** Pie chart showing the proportion of cells labeled with GFP alone, tdTomato alone, or both. Labeled cells were counted in the M1 region (542.5 µm × 722.4 µm) (*n* = 5 mice, 1 section/mouse). A total of 2,349 fluorescently labeled cells were counted, among which 1,094 were GFP-positive only, 1,207 were tdTomato-positive only, and 48 were co-labeled with both GFP and tdTomato.

Supplementary figure 7

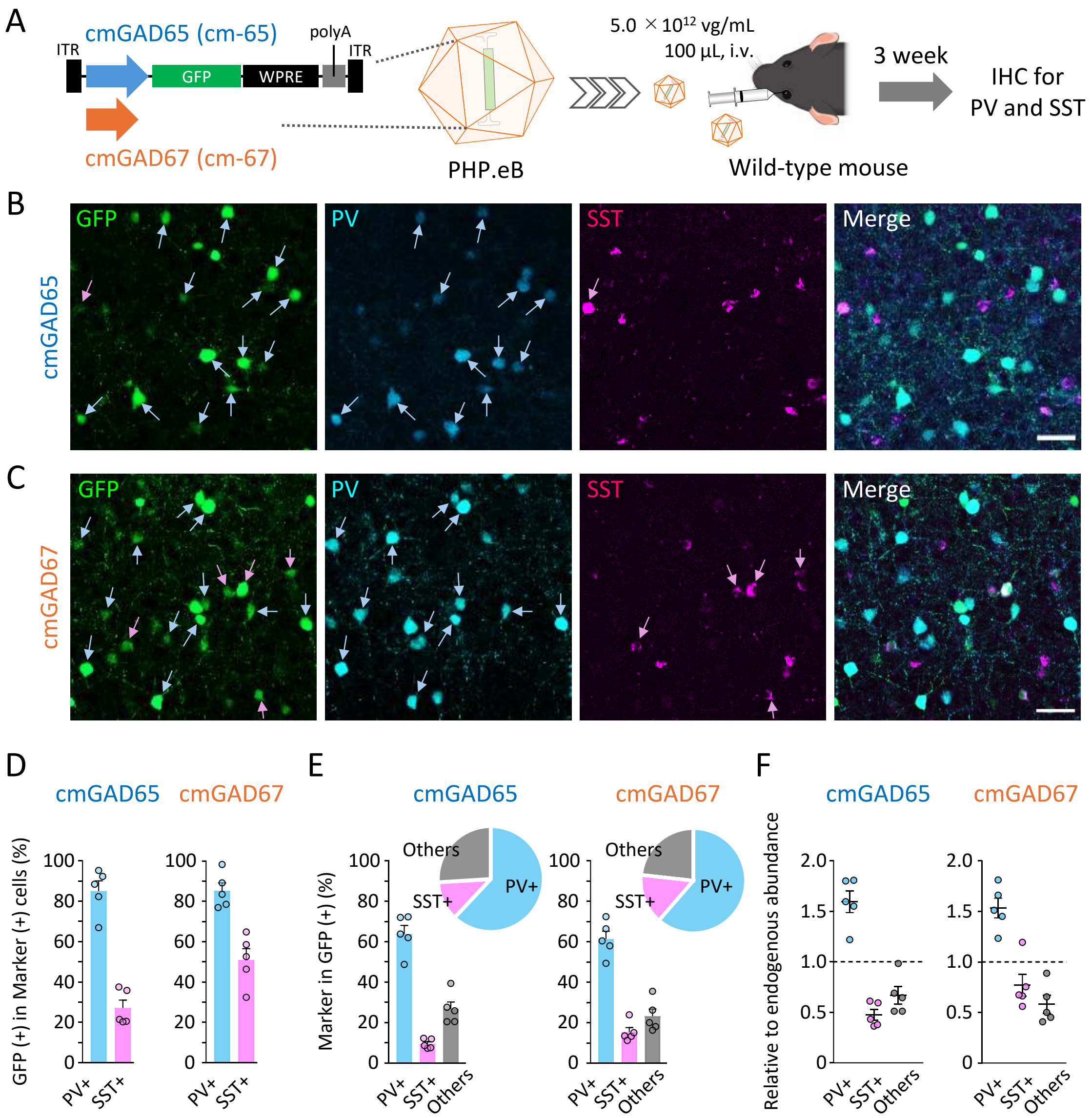

**Supplementary figure 7. Subtype specificity of cmGAD65 and cmGAD67 promoters in PV and SST interneurons.**

**A**. Schematic of the experimental procedure. AAV-PHP.eB vectors expressing GFP under the control of the cmGAD65 or cmGAD67 promoter were intravenously injected into wild-type mice. Three weeks later, sagittal brain sections were prepared and subjected to immunohistochemistry (IHC).

**B**, **C**. Immunohistochemical staining of the motor cortex for parvalbumin (PV, blue channel) and somatostatin (SST, magenta channel) in mice injected with PHP.eB carrying the cmGAD65 promoter (B) or cmGAD67 promoter (C). Cyan arrows indicate PV-positive ($PV^+$) neurons, and magenta arrows indicate SST-positive ($SST^+$) neurons. Scale bars, 50 µm.

**D**. Quantification of the proportion of $PV^+$ and $SST^+$ neurons labeled with GFP. Each dot represents data from an individual mouse.

**E**. Bar graphs and pie charts showing the composition of $PV^+$, $SST^+$, and double-negative cells among GFP-expressing neurons.

**F**. Relative tropism of the cmGAD promoter-driven AAV-PHP.eB vectors for different GABAergic neuron subtypes ($PV^+$, $SST^+$, or others) in the mouse cortex. In the mouse cortex, $PV^+$ neurons account for 40%, $SST^+$ neurons for 15%, and other subtypes for 45% of all GABAergic neurons [1]. The ratio of each GFP-expressing subtype (from E) was divided by its expected proportion in the cortex to calculate relative enrichment. A value near 1 indicates no subtype preference, whereas a value greater than 1.5 for $PV^+$ neurons indicates selective gene expression in this subtype.

Supplementary figure 8

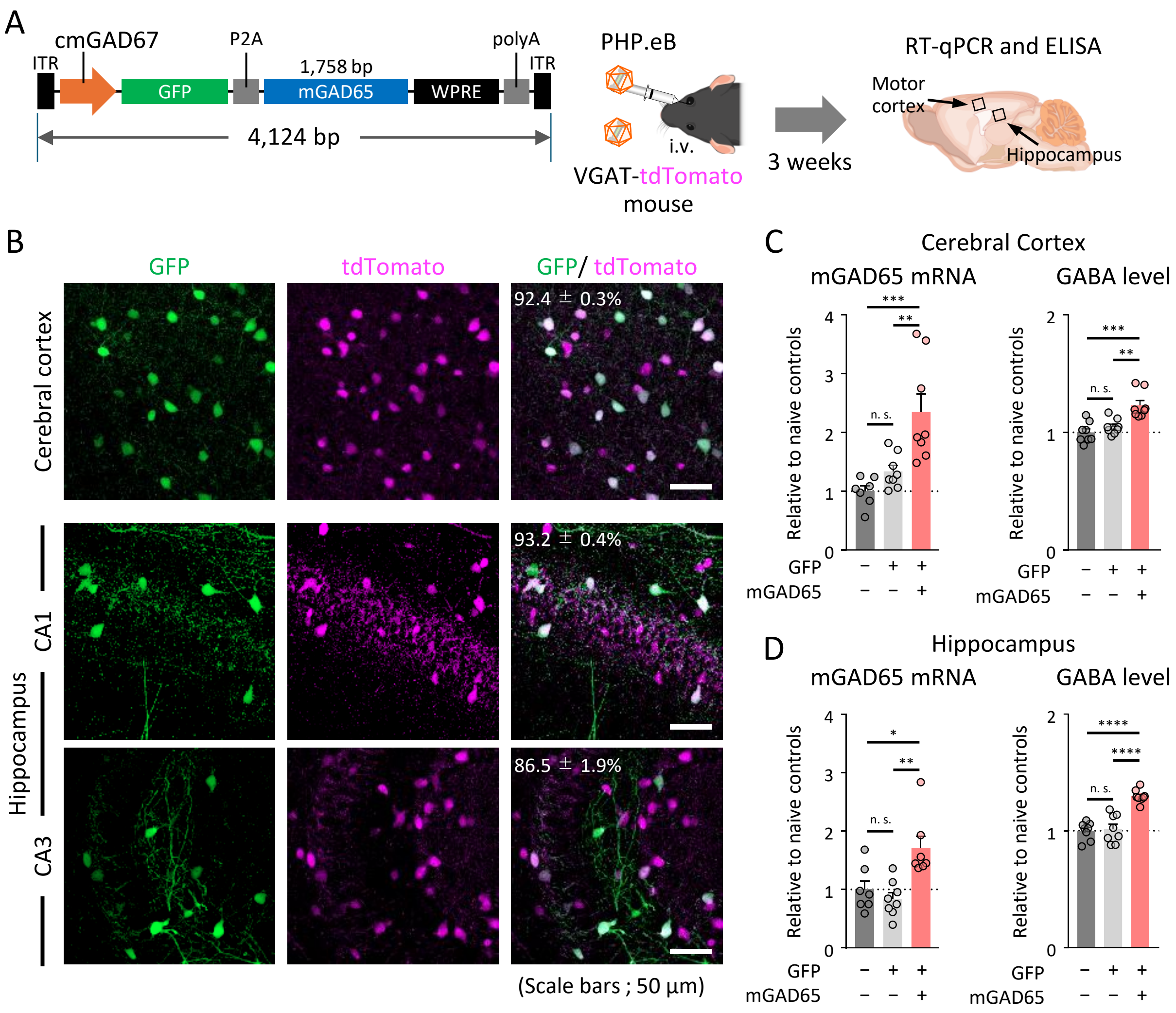


**Supplementary figure 8. AAV-cmGAD67-GAD65 elevates cortical and hippocampal GABA levels.**

**A.** Schematic representation of the experimental procedure. AAV-PHP.eB expressing GFP and *mGAD65* under the control of the cmGAD67 promoter was intravenously administered to *VGAT*-tdTomato mice. Three weeks post-injection, tissue samples from the primary motor cortex and hippocampus were collected for quantification of *mGAD65* mRNA using RT-qPCR and measurement of GABA levels via ELISA.

**B.** Fluorescence images of native GFP and native tdTomato in the primary motor cortex and hippocampal CA1 and CA3 regions. The numbers in the upper left corner of the rightmost panel indicate the percentage of tdTomato-positive (inhibitory) neurons among GFP (*mGAD65*)-expressing cells (Average ± SEM., $n = 5$ mice). Scale bars, 50 µm.

**C, D.** Quantification of *mGAD65* mRNA (left) and GABA levels (right) in the primary motor cortex (**C**) and hippocampus (**D**). The values were normalized to those obtained from untreated mice (leftmost bar). Relative expression levels are shown for mice injected with AAV-PHP.eB expressing GFP alone and those expressing both GFP and *mGAD65*. Dots in the graphs represent individual mouse data points. *$p < 0.05$, **$p < 0.01$, ***$p < 0.001$, ****$p < 0.0001$, n.s., not significant; one-way ANOVA with Tukey's post hoc test.

Supplementary figure 9

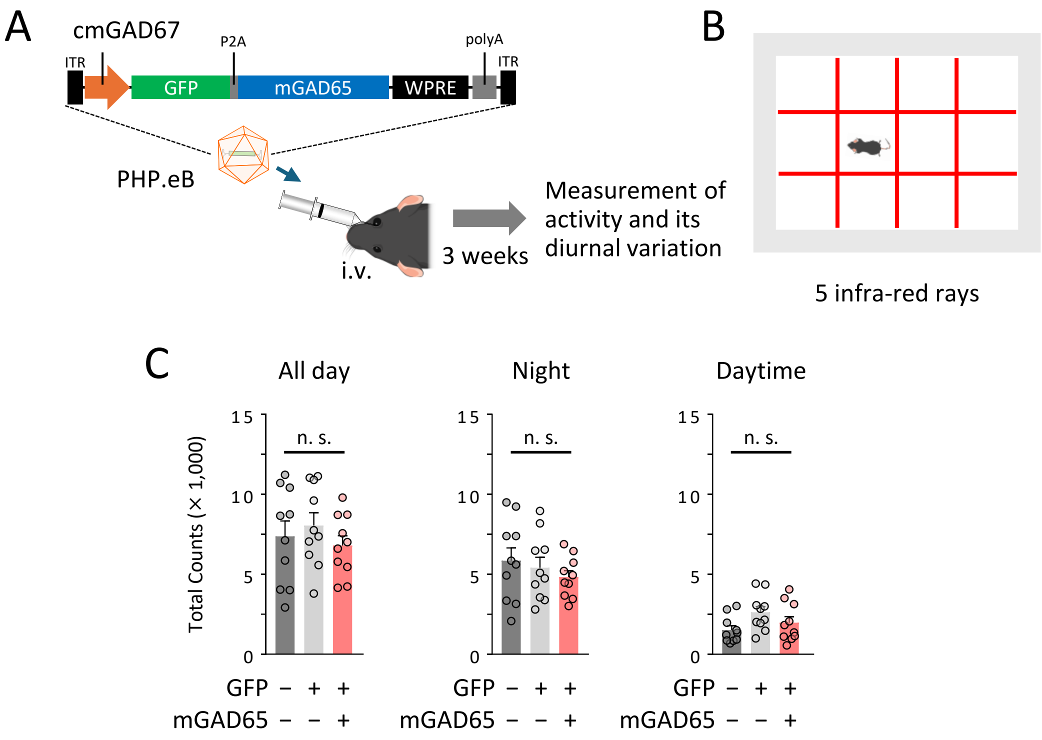


**Supplementary figure 9. Safety assessment: AAV-cmGAD67-GAD65 does not alter circadian activity.**

**A.** Schematic representation of the experimental procedure. AAV-PHP.eB expressing GFP and GAD65 under the control of the cmGAD67 promoter was intravenously administered. Three weeks post-injection, locomotor activity and its circadian variations were measured.

**B.** Illustration of the activity monitoring apparatus. Mice were placed in a cage segmented by five horizontal and five vertical infrared beams, and locomotor activity was quantified by counting the number of infrared beam crossings.

**C.** Quantification of the number of infrared beam crossings over the entire 24-hour period, during the nocturnal phase, and during the diurnal phase. Dots in the graph represent individual mouse data points. n.s., not significant; one-way ANOVA with Tukey's post hoc test.

### Supplementary figure 10

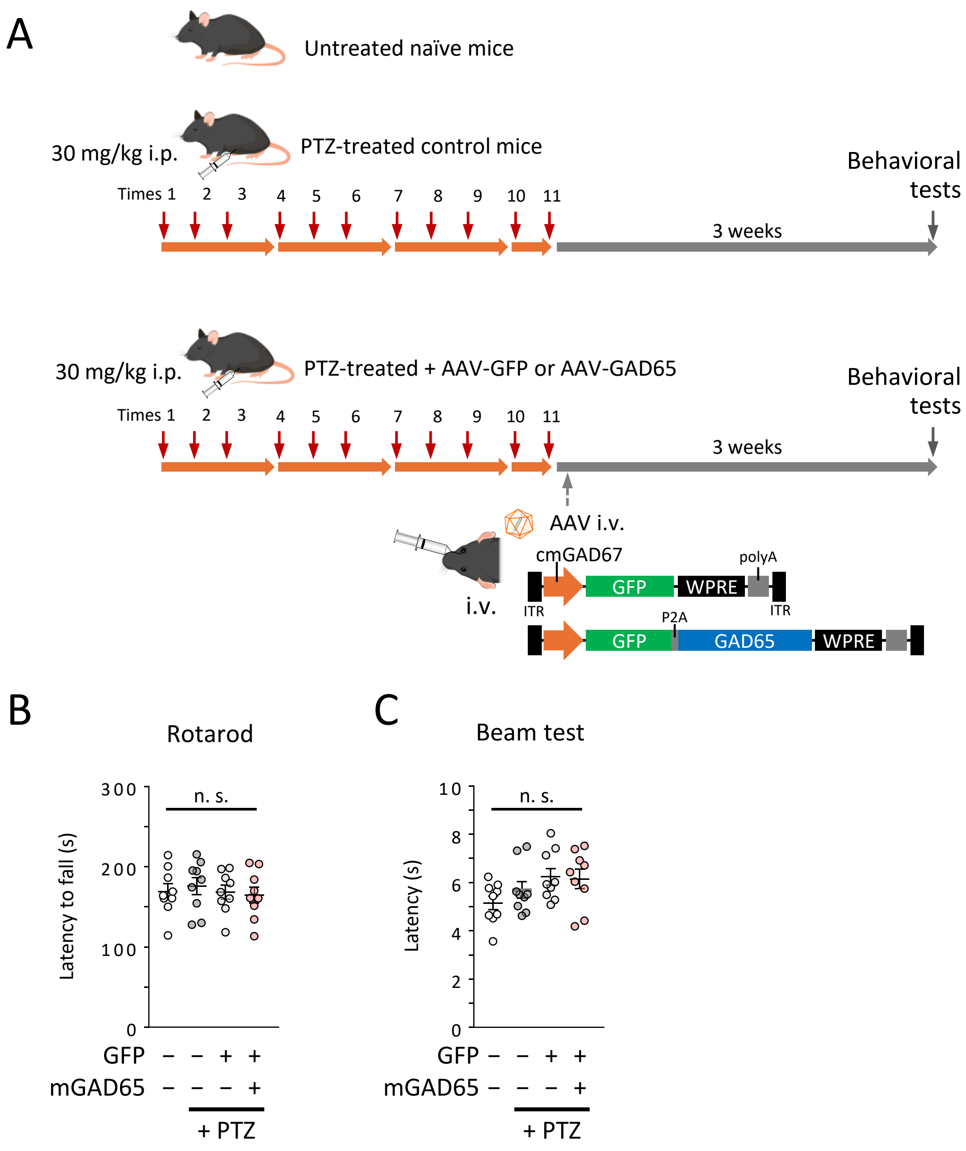


**Supplementary figure 10. Safety assessment: AAV-cmGAD67-GAD65 does not impair motor coordination.**

**A.** Schematic representation of the experimental procedure. Naïve mice were used as controls. The rest of three groups received pentylenetetrazole (PTZ, 30 mg/kg) intraperitoneally three times per week (every other day) for a total of 11 sessions. One group were treated only with PTZ. Two other groups received intravenous injection of AAV-PHP.eB expressing GFP-P2A-GAD65 or GFP alone under the control of the cmGAD67 promoter. Behavioral experiments were conducted three weeks later.

**B.** Results of the rotarod test. The latency to fall from the rotating rod was measured.

**C.** Results of the beam test. Mice were placed at one end of a narrow metal beam, and the time taken to traverse the beam and enter a shelter at the opposite end was recorded. Dots in the graphs represent individual mouse data points. n.s., not significant; ANOVA with Tukey's post hoc test.

Supplementary figure 11

#### Statics for Main Figures

| | |
|---|---|
| Figure 1 | C, D: Unpaired t-test |
| Figure 3 | D: Mann-Whitney tests<br>G: Paired t-test |
| Figure 4 | B: Pearson correlation<br>D(b, c), E(b,c), F(b,c): Paired t-test<br>G, H: one-way ANOVA Turkey's post hoc test<br>I: Energy distance test for equality of distributions |
| Figure 5 | B, C: one-way ANOVA Turkey's post hoc test |
| Figure 7 | B, E: two-way ANOVA repeated measures (Group effect)<br>C, F: Gehan-Breslow-Wilcoxon test |

#### Statics for Supplementary Figures

| | |
|---|---|
| Figure 1 | F, H: one-way ANOVA Dunnett's post hoc test |
| Figure 2 | C, E: one-way ANOVA Dunnett's post hoc test |
| Figure 3 | C, D: one-way ANOVA Dunnett's post hoc test |
| Figure 4 | E, F, G: one-way ANOVA Dunnett's post hoc test |
| Figure 5 | B: one-way ANOVA Dunnett's post hoc test |
| Figure 8 | C, D: one-way ANOVA Turkey's post hoc test |
| Figure 9 | C: one-way ANOVA Turkey's post hoc test |
| Figure 10 | B, C: one-way ANOVA Turkey's post hoc test |

Supplementary methods

**Promoter construct generation**

Stepwise truncations and deletions of the *Gad1* locus were performed to identify regulatory regions including Ri, Rii, and ETSS. Constructs were cloned into AAV-PHP.eB backbones driving GFP expression for in vivo analysis (Supplementary figure 1-3).

**RT-qPCR quantification**

Total RNA was extracted from dissected cortical tissue, reverse transcribed. GFP (Supplementary figure 5) and mGAD65 (Supplementary figure 8) expressions in mice brain at 3 weeks after AAV vector administration was analyzed by quantitative RT-PCR. Cortical or hippocampal samples were homogenized in TRizol Reagent (Thermo Fisher Scientific) on ice and cooled in liquid nitrogen. RNA was extracted by RNeasy Lipid Tissue Mini Kit (Qiagen, Hilden, Germany) and cDNA was prepared by ReverTra Ace reverse transcriptase (Toyobo, Osaka, Japan). GFP expression was determined by quantitative real-time PCR (TP900; TaKaRa Bio, Shiga, Japan) using Power SYBR Green Master Mix (Thermo Fisher Scientific,), mGAD65 expression was determined by quantitative real-time PCR (T900 or TP970; TaKaRa Bio) using THUNDERBIRD SYBR qPCR Mix (Toyobo). Relative expression levels were normalized to GAPDH or β-actin.

The primers used for supplementary figure 5 are as follows.
GFP: 5′-GGA CGA GCT GTA CAA GTA AAG-3′ (Forward) and 5′-GGG AAG CAA TAG CAT GAT ACA AAG G-3′ (Reverse)
GAPDH: 5'-GTG TTC CTA CCC CCA ATG TG-3' (Forward) and 5'-GGT GGA AGA GTG GGA GTT GCT G-3' (Reverse)

The primers used for supplementary figure 8 are as follows.
mGAD65: 5′-ACT ACT GGG TTT GAG GCA CA-3′ (Forward) and 5′-TGC GCA AAC TAG GAG GTA CA-3′ (Reverse)
β-actin: 5′-ACC AAC TGG GAC GAT ATG GAG AAG A-3′ (Forward) and 5′-TAC GAC CAG AGG CAT ACA GGG ACA A-3′ (Reverse)

**Immunohistochemistry**

Supplementary figure 7 related to Figure 2. The same antibody used in Figure 2 was used. Co-localization of GFP with $PV^+$ and $SST^+$ cells was analyzed by BZ-800 and confocal microscopy.

**GABA quantification**

Three weeks after injection of AAV vector or PBS, cortical tissue (near M1) and hippocampal samples were collected. GABA concentrations were quantified using a Mouse GABA enzyme-linked immuno sorbent assay (ELISA) Kit (ab287793; Abcam) according to the manufacturer's instructions. Brain tissue was homogenized in PBS, and total protein concentrations were measured using the Micro BCA™ Protein Assay Kit (Thermo Fisher Scientific). Samples were diluted to equal protein concentrations prior to ELISA. Absorbance values were measured using an Infinite M Nano microplate reader (Tecan, Männedorf, Switzerland). Values were normalized to protein concentration in each sample.

**Behavioral paradigms**

**Locomotor Activity Monitoring:**

Infrared beam breaks were recorded over a 24-hour period using the Locomotor Activity Measuring System (LS-5; MELQUEST, Tokyo, Japan) to assess circadian activity. Mouse activity was monitored continuously throughout the day, starting at 12:00 noon, using infrared light emitted from the device.

Supplementary methods

**Rotarod and Beam Tests:**

Motor coordination was evaluated using the rotarod latency to fall and beam traversal time based on previous report [2]. Three weeks after AAV or PBS administration, rotarod performance was assessed using a treadmill device (MK-610A/RKZ; Muromachi Kikai). Each mouse was tested three times at 30-minute intervals. The rotational speed was gradually increased from 4 rpm to 50 rpm over 300 seconds, and the time to fall from the rod was recorded. For the beam test, mice were placed at one end of a narrow bar (800 mm long, 11 mm in diameter) and allowed to walk toward a goal box located at the opposite end. This task was repeated three times, and the average time required to traverse the beam was recorded as a measure of motor coordination.
